# Pharmacologic eIF2B Activation Rescues Neuropathy in CMT2 Subtypes by Normalizing the Integrated Stress Response

**DOI:** 10.64898/2026.08.05.743063

**Authors:** Mary McMahon, Steve Lianoglou, Sridhar Narayan, Jiacheng Zhang, Shilin Chen, Jinxing Li, William Kluwe, Yang Liu, Bin Cao, Jie Luo, Jijun Chen, Xiaojin Zhang, Sizhu Lu, Melanie Das, Anil C. Nair, Xinru Meng, Litao Sun, David Gong, Mona M Freidin, Charles K Abrams, Yang Li, Peng Yue, Paul R. August

**Affiliations:** ReviR Therapeutics Inc., Brisbane, California, USA; Xili Technology (Shenzhen) Co., Ltd., Shenzhen, Guangdong, China; XtalPi Inc., 207 Huanqiao Rd, Pudong New Area, Shanghai, China; Department of Pharmacology and Toxicology, University of Arizona, Tucson AZ 85721; Shenzhen Key Laboratory of Pathogenic Microbes and Biosafety, Shenzhen Campus of Sun Yat-sen University, Sun Yat-sen University, Shenzhen 518107, China; Department of Neurology and Rehabilitation, University of Illinois Chicago, College of Medicine, Chicago, IL, USA

## Abstract

Among the many subtypes of Charcot–Marie–Tooth (CMT) disease, several result from mutations in genes encoding aminoacyl-tRNA synthetases, enzymes required for tRNA charging during cytoplasmic and mitochondrial translation. We report that activation of the integrated stress response (ISR) pathway is a shared molecular feature of tRNA synthetase-associated and other axonal CMT subtypes. RTX-117, a CNS-penetrant small molecule currently in Phase 1 clinical trials, targets eukaryotic initiation factor 2B (eIF2B), a key modulator of protein synthesis and the ISR pathway. Using cryo-EM studies, we have characterized the binding mode of RTX-117 to the eIF2B decamer. In *Gars*^P278KY/+^ mice, which develop early onset motor defects and axonal pathology that recapitulate CMT2D symptoms from tRNA synthetase mutations, RTX-117 treatment started after disease onset reduced chronic ISR activation and produced significant functional and electrophysiological improvement. We further identify ISR targets, including secreted proteins such as GDF15 and FGF21 that may serve as translational biomarkers for treatment response to RTX-117 in CMT disease. Broader surveillance of the ISR pathway across models of neurodegeneration reveals strong activation in several diseases and a correlation with disease progression, particularly in models of Alzheimer’s disease. These findings identify chronic ISR activation as a recurrent, though not universal, pathological mechanism of neurodegenerative disease models. Overall, our study identifies candidate biomarkers for CMT disease subtypes associated with defects in translational homeostasis and supports eIF2ɑ-ATF4 axis modulation as a promising therapeutic strategy for this disease class.

**One Sentence Summary:** RTX-117, a clinical stage eIF2B activator, blunts chronic ISR activation and improves nerve and motor function in a mouse model of Charcot-Marie-Tooth Disease Type 2D.

## INTRODUCTION

Charcot-Marie-Tooth (CMT) disease is a genetically and clinically heterogeneous group of inherited peripheral neuropathies, characterized by progressive distal muscle atrophy and sensory loss(*1–3*). Mutations in more than 130 genes cause CMT disease, classified primarily into demyelinating (CMT type 1, CMT1), axonal (CMT type 2, CMT2), and intermediate (CMTI) forms(*4–5*). CMT1, the largest subtype, is driven predominantly by mutations in genes involved in myelination. CMT2-associated mutations, in contrast, disrupt axonal transport, mitochondrial dynamics, tRNA charging, and proteostasis, placing motor and sensory neurons under sustained metabolic and proteotoxic stress(*6–8*). Clinically, CMT2 causes progressive distal muscle atrophy and weakness, sensory loss, and reduced compound muscle action potential amplitudes reflecting primary axonal pathology rather than demyelination. Onset typically occurs in the first or second decade of life, though severity and progression vary considerably among subtypes and individuals(*9–10*). Over time, affected individuals develop foot deformities, gait disturbances, and loss of fine motor function. No intervention currently halts or reverses these symptoms. The length and metabolic demands of peripheral axons likely render them particularly vulnerable to proteotoxic stress, which must be managed over millimeter-to-meter distances.

The heterogeneity of CMT disease subtypes has historically hindered therapeutic development, with no disease-modifying therapies approved to date(*11*). Targeting individual gene defects is unlikely to yield broadly effective treatments, because each therapy would only benefit a limited subset of patients. Instead, strategies that target shared pathogenic mechanisms, such as mitochondrial dysfunction and impaired proteostasis, offer a tractable path to a disease-modifying approach spanning genetically diverse CMT subtypes. The largest gene family implicated in CMT2 encodes enzymes known as aminoacyl-tRNA synthetase (ARS), with mutations in up to 8 members (AARS1, DARS2, GARS1, HARS1, KARS1, MARS1, WARS1, and YARS1) identified in CMT2 subtypes to date(*12–16*). Although genetically diverse, ARS-associated CMT2 forms a mechanistically unified disease class. These enzymes catalyze cytoplasmic and mitochondrial tRNA charging with their cognate amino acids, a key step in protein synthesis and cellular homeostasis(*17*), potentially highlighting a common molecular pathology shared across major CMT disease subtypes.

The integrated stress response (ISR) pathway is a conserved translational regulatory program activated by numerous forms of cellular stress and controlled by stress sensing kinases (*18-19*). In several model organisms, ARS-associated CMT2 mutations activate the stress sensing protein kinase General Control Nonderepressible 2 (GCN2), leading to phosphorylation of the alpha subunit of eukaryotic initiation factor 2 (eIF2), a key regulator of translation initiation(*20–23*). Phosphorylation of eIF2α inhibits the guanine nucleotide exchange factor (GEF) activity of eukaryotic Initiation Factor 2B (eIF2B), suppressing cap-dependent translation while selectively enhancing translation of mRNAs encoding stress-responsive proteins including the transcription factor ATF4. Both genetic and pharmacological inhibition of GCN2 reduces chronic ISR activation and rescues neuropathy in CMT2D mouse models driven by GARS1 mutations(*23*). The same ISR mechanism extends to a broader class of disorders affecting mitochondrial tRNAs, suggesting shared pathogenic mechanisms across metabolic, neurological, and neuromuscular phenotypes(*24*). While transient ISR pathway activation is adaptive, chronic engagement impairs proteostasis, disrupts mitochondrial function, and drives progressive neuronal dysfunction across a number of neurodegenerative diseases(*18-19*). In Vanishing White Matter (VWM) disease, mutations in eIF2B cause chronic ISR activation and progressive white matter degeneration thereby establishing eIF2B dysfunction as a driver of neurodegeneration in humans(*25*). These observations provide genetic validation of the eIF2-eIF2B axis as a central regulator of neuronal homeostasis and implicate chronic ISR dysregulation as a possible pathogenic and therapeutically tractable axis in CMT2.

Pharmacological modulation of the ISR has been pursued through multiple strategies, including inhibition of upstream stress sensing kinases such as PERK, GCN2, HRI, and PKR. However, a narrow therapeutic window has been reported for several of these kinases; for example, dose-limiting pancreatic toxicity has been reported with PERK inhibitors(*26*). Furthermore, each kinase responds to distinct upstream signals and engages signaling networks beyond eIF2α phosphorylation, complicating therapeutic specificity(*26–27*). In contrast, direct activation of eIF2B offers a mechanistically distinct approach by restoring GEF activity downstream of eIF2α phosphorylation, thus preserving upstream stress sensing mechanisms. Small molecules that bind and stabilize the active (A-state) conformation of eIF2B such as ISRIB, restore protein synthesis and demonstrate efficacy across multiple models of neurodegeneration and injury, validating eIF2B as a key therapeutic target within the ISR pathway(*28–33*). Clinical-stage eIF2B activators, such as DNL343 and fosigotifator, have advanced into trials for ALS and VWM disease and appear well tolerated in humans(*34–35*). Building on this rationale, we developed a small molecule eIF2B activator with greater potency and improved pharmacological properties compared with earlier eIF2B activators, designed to enable sustained and controllable restoration of translational homeostasis. In this study, we report that the investigational compound RTX-117, currently in a Phase 1 trial in healthy participants (<u>ChiCTR2600118758</u>), stabilizes the eIF2B complex and attenuates ATF4-driven stress signaling. We demonstrate a time- and dose-dependent recovery of motor axon function with RTX-117 treatment in a mouse model of CMT2D, consistent with a dose-dependent ISR pathway suppression, supporting eIF2B activation as an effective *in vivo* modifier of ISR-driven peripheral neuropathy. Our findings point to pharmacologic eIF2B activation as a viable disease-modifying strategy for CMT2 subtypes and positions ISR normalization as a scalable therapeutic entry point for axonal neuropathies and potentially other diseases driven by chronic translational stress.

## RESULTS

### Discovery of RTX-117, a small molecule eIF2B activator with translational pharmacology

Given the need for sustained ISR normalization in chronic neurodegenerative conditions, we hypothesized that a next generation eIF2B activator with improved potency and optimized pharmacokinetic (PK) properties would be required to achieve consistent target engagement in the nervous system. To design a molecule having these characteristics, we built a computational model based on the published structure of ISRIB-bound eIF2B(*36*). Using a variety of computer-aided drug design (CADD) tools including XPOSE and XFEP, a series of compounds were designed and synthesized. An ATF4-luciferase reporter system was used as the primary assay, while in vitro absorption, distribution, metabolism, and excretion (ADME) assays were used to optimize drug metabolism and pharmacokinetics (DMPK). These efforts led to the identification of RTX-117 as a brain-penetrant molecule suitable for once-a-day oral dosing (**Fig. 1A and Table S1**). The full details of the medicinal chemistry and structure-activity relationship (SAR) optimization will be reported elsewhere. The binding affinity of RTX-117 to eIF2B was quantified using a fluorescence polarization competition assay, in which unlabeled RTX-117 displaced FAM-labeled RTX-117 bound to the eIF2B decamer with an IC_50_ of 41.8 nM (**Fig. 1B**). To define the molecular basis of eIF2B binding by RTX-117, we solved a cryogenic electron microscopy (cryo-EM) structure of RTX-117 bound to the human eIF2B decamer. The final electron density map, derived from 263,354 particles, achieved an average resolution of 2.8 Å with C1 symmetry. The overall resolution of the map ranged from 2.5 to 4.5 Å, while the local resolution of the binding site showed that the density for RTX-117 was between 2.5 to 3 Å (**Fig. 1C**). The structural model was built by fitting the density map to a previously reported model of eIF2B bound to ISRIB(*36*).

**Figure 1.**
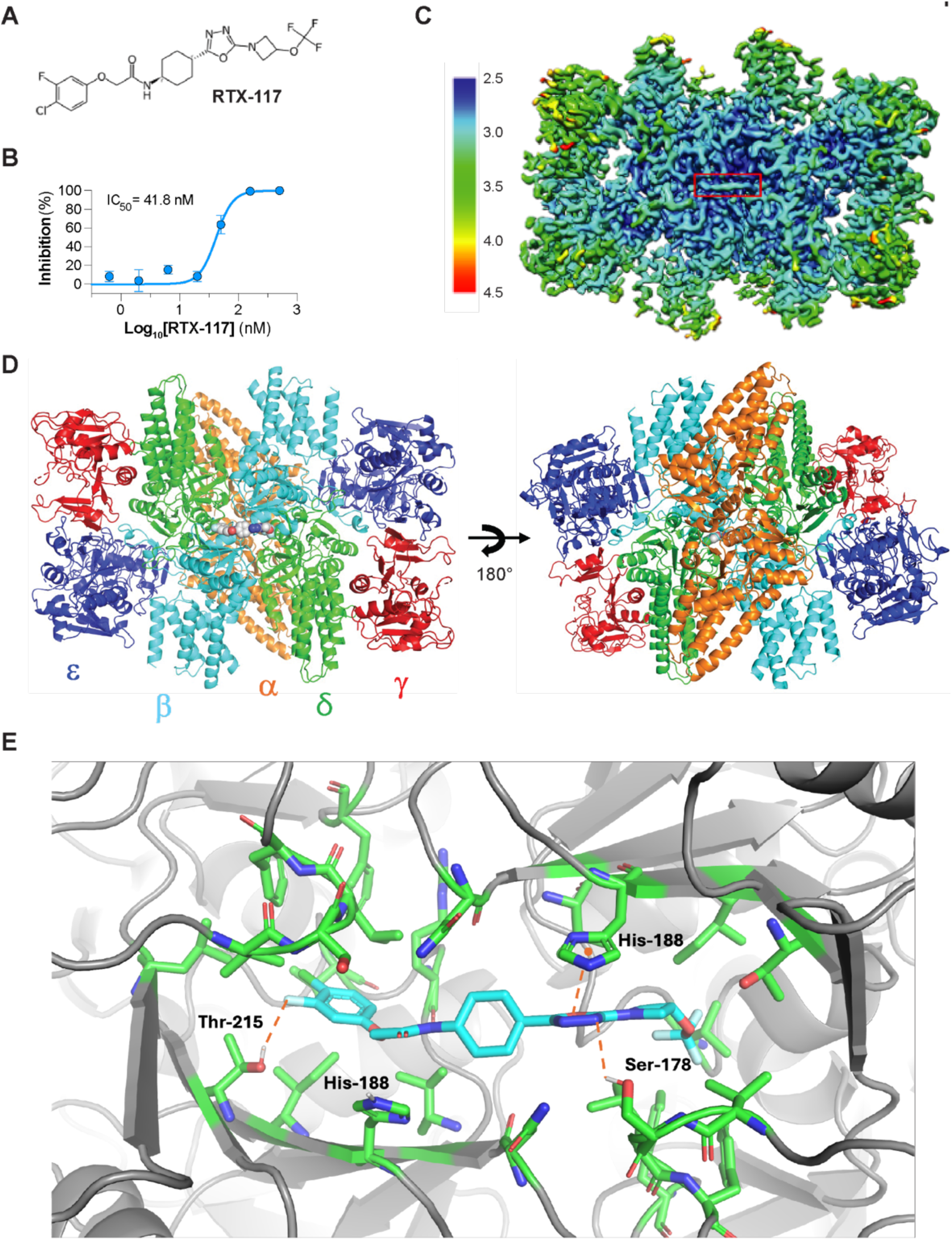
Biophysical and structural analysis of RTX-117 binding to eIF2B. (A) Chemical structure of RTX-117 (C_20_H_21_ClF_4_N_4_O_4_). (B) Fluorescence polarization competition assay showing displacement of FAM-labeled RTX-117 from the eIF2B decamer by increasing concentration of unlabeled RTX-117. Data represent mean ± SD. from at least three independent replicates. (C) Cryo-electron microscopy (cryo-EM) density map of RTX-117 bound to the human eIF2B decamer. Local resolution varies across structure, with the ligand-binding region resolved at 2.5 to 3.0 Å. The RTX-117 binding site is highlighted with a red box. (D) Ribbon representation of the global structure of the eIF2B decamer bound to RTX-117, shown in two orientations. Individual subunits are colored to illustrate the overall architecture and ligand positioning within the complex. (E) Close-up view of the RTX-117 binding pocket within eIF2B, highlighting important residues and intermolecular interactions that contribute to ligand binding and stabilization of the complex. Red dotted lines highlight key interactions.

RTX-117 binds at the canonical ISRIB pocket at the interface of eIF2B subunits, where it stabilizes the decameric complex (**Fig. 1D**). The binding site is flanked by two monomers each of the α, β, and δ subunits, and the overall binding mode is similar to that of ISRIB, where the bound ligand brings together and stabilizes the eIF2B decamer. In addition to hydrophobic interactions characteristic of ISRIB, RTX-117 forms novel polar contacts with Thr-215 and Ser-178 and engages in π-stacking with His-188 (**Fig. 1E**). These additional interactions likely contribute to the enhanced affinity and potency of RTX-117 over ISRIB and other eIF2B activators. Comparison of the RTX-117-bound and apo eIF2B structures reveals minimal structural rearrangement, consistent with a mechanism in which RTX-117 stabilizes the active conformation of eIF2B rather than inducing a distinct conformational change (**Fig. S1A**). The ligand interaction map further highlights the network of hydrophobic and polar contacts that underlie RTX-117 binding specificity and affinity (**Fig. S1B**). These data establish RTX-117 as a high-affinity eIF2B activator that stabilizes the active conformation of the complex and supports restoration of translational control upon ISR activation.

### RTX-117 suppresses stress-induced ATF4 signaling and normalizes ISR pathway activation in human cells

Having established the structural basis of RTX-117 binding to eIF2B, we next evaluated whether enhanced eIF2B binding and decamer stabilization produced functional suppression of ISR signaling in human cells. First, we assessed whether RTX-117 modulated ATF4-driven ISR signaling in HEK293T cells using an established ATF4-luciferase reporter assay(*28*). Treatment with thapsigargin, an inhibitor of the sarco/endoplasmic reticulum Ca^2+^-ATPase (SERCA) pump that induces endoplasmic reticulum (ER) stress, robustly activated the reporter within 6 hours. Co-treatment with RTX-117 suppressed ATF4-luciferase reporter activity in a concentration-dependent manner, demonstrating potent inhibition at low nanomolar concentrations (IC_50_ = 1.59 nM) **(Fig. 2A)**. RTX-117 was more than 10-fold more potent than DNL343(*32, 34*) in this assay (**Fig. S2A**). Suppression of endogenous ATF4 protein levels following thapsigargin treatment was confirmed by western blot (**Fig. S2B**).

**Figure 2.**
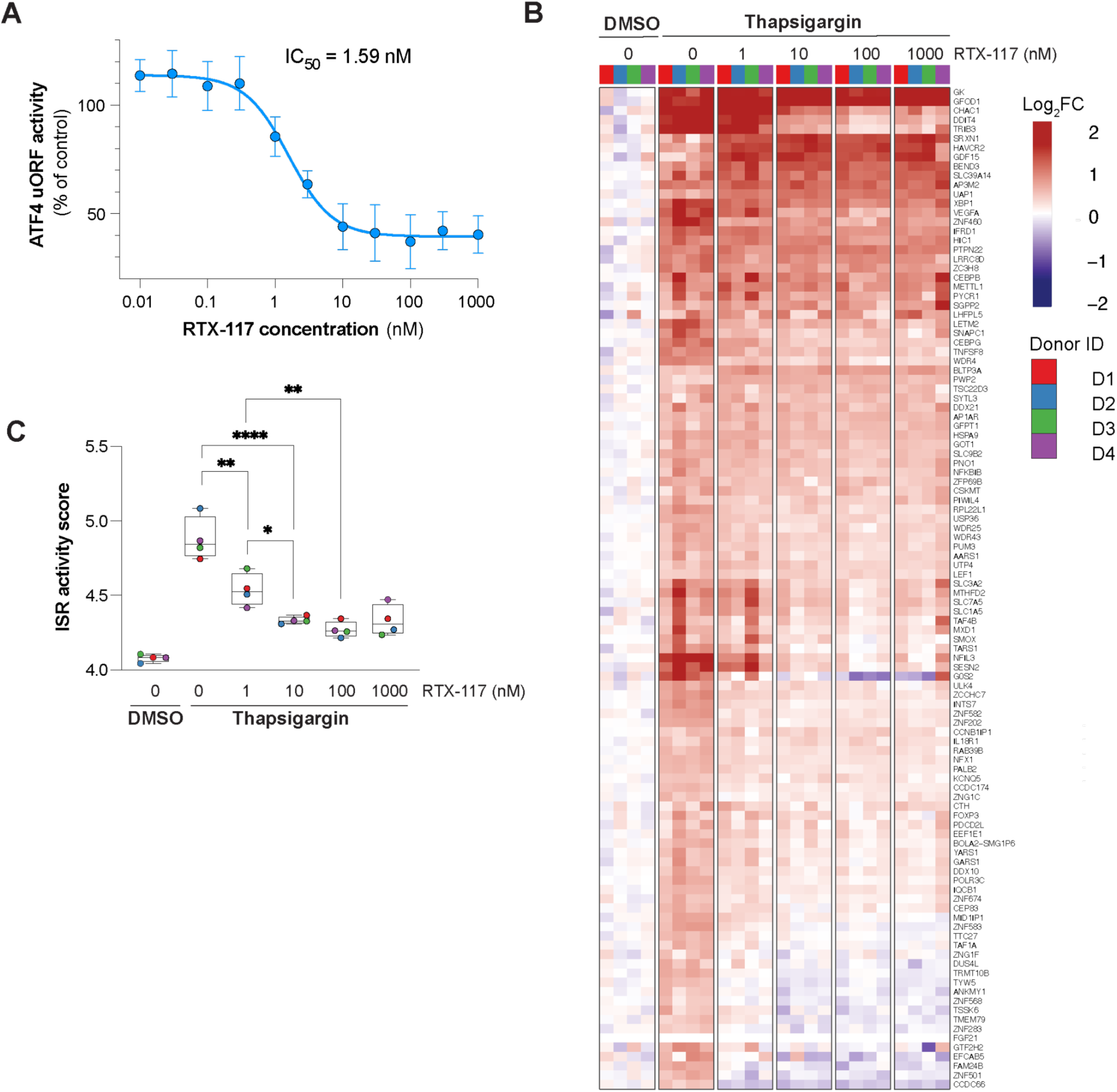
RTX-117 suppresses ATF4 activation and integrated stress response (ISR) signaling in human cells. (A) Concentration-dependent inhibition of ATF4-luciferase reporter activity in HEK293T cells stimulated with thapsigargin. Cells were treated with increasing concentrations of RTX-117, and ATF4 translation was quantified using a reporter assay (n=5 experiments). Data are presented as mean ± SD. Nonlinear regression analysis yielded an IC_50_ of 1.59 nM. (B) Heatmap visualization of thapsigargin-induced ISR modulation by RTX-117. Rows represent the 106-gene subset of the ISR Class 1 signature that is upregulated by thapsigargin and columns represent samples from individual donors across conditions. Color scale represents log_2_ fold change for each gene per sample as compared to its mean expression across the DMSO control samples. Expression values are first donor-corrected prior to DMSO normalization. Although GDF15 and FGF21 are not part of the thapsigargin-induced signature, they have been added here due to their importance as potential translational biomarkers. (C) Single-sample ISR activity scores calculated from the genes in the heatmap using their eigen-weighted mean (see methods). Points represent individual donor samples and boxes indicate median and interquartile range. Brackets indicate comparisons between indicated treatment groups. Stars indicate statistical significance from Holm’s adjusted p-values (see methods). *P < 0.05, **P < 0.01, ****P < 0.0001.

We next evaluated whether these effects extended to primary human cells. Peripheral blood mononuclear cells (PBMCs) from four independent donors were stimulated with thapsigargin and analyzed by bulk RNA-seq to assess ISR pathway activation. Thapsigargin broadly dysregulated the transcriptome with approximately 3,000 genes up- and down-regulated by more than 30% at an FDR < 5% (**Fig. S3A**). The transcriptional response to ISR activation varies across cell types and depends on the underlying trigger, though it consistently engages genes involved in translation, amino acid metabolism, redox homeostasis, apoptosis, and proteostasis(*37*). To identify ISR activation from gene expression data, we curated a set of seven primary ISR transcriptional signatures from the literature, each consisting of genes that are either up-regulated during ISR activation or inferred to be direct targets of ATF4 (see the “ISR Signature Collection” methods section, and Table S2). We performed gene set enrichment analysis (GSEA)(*38, 39*) with this signature collection to validate ISR activation by thapsigargin (**Fig. S3B-C and Table S3A**). To better quantify the degree of ISR modulation by RTX-117 in this context, we identified the subset of 106 ISR Class 1 genes that are upregulated under thapsigargin treatment (**Fig. S3D and Table S3B**) and tracked their differential expression with increasing doses of RTX-117. This gene set was further used for visualization and statistical analysis.

Co-treatment with RTX-117 attenuated ISR pathway activation across all four donors, with particularly strong suppression of CHAC1, DDIT4, and TRIB3, among several other ISR Class 1 genes, in a concentration-dependent manner **(Fig. 2B and Fig. S3E-G).** To quantify ISR modulation more accurately at the pathway-level, we computed a single-sample ISR activity score using the 106-gene Class 1 signature (see methods section). Thapsigargin elevated the ISR activity scores relative to vehicle controls, while RTX-117 reduced this response in a concentration-dependent manner across donors (**Fig. 2C and Table S3A**), with reductions in ISR pathway activation observed at concentrations as low as 1 nM. These findings demonstrate that RTX-117 is a potent eIF2B activator capable of suppressing stress-induced ATF4 upregulation and normalizing the downstream ISR transcriptional program in primary human cells.

### RTX-117 reduces ISR and improves neuropathology in a mouse model of VWM disease

To establish target engagement of RTX-117 *in vivo,* we turned to a well-characterized mouse model of VWM disease and ISR activation caused by destabilizing mutations in the eIF2B complex. *Eif2b5*^R191H/R191H^ knock-in (hereafter referred to as *Eif2b5* KI) mice, which carry the pathogenic allele orthologous to the human eIF2B5 R195H mutation that causes VWM disease, develop myelin loss, progressive motor dysfunction, and chronic ISR activation in the central nervous system(*31, 32*). *Eif2b5* KI mice were treated orally once daily with RTX-117 (0.1–10 mg/kg) for up to six months and monitored for disease progression and ISR activation (**Fig. S4A**). PK analysis confirmed systemic exposure and brain penetration of RTX-117 across all dose groups, with comparable unbound compound levels detected in both plasma and brain (**Fig. S4B**). Consistent with previous reports, transcriptomic analysis of brain tissue from vehicle-treated *Eif2b5* KI mice revealed ISR pathway activation relative to WT controls (**Fig. S4C** and **Table S3C)**. A 37-gene signature was generated from the genes up-regulated in the ISR Class 1 signature (> 30% increase at FDR < 5%) and was used to assess pharmacological reduction of ISR activity (**Table S4D**). RTX-117 treatment normalized ISR pathway activation in a dose-dependent manner, with reductions in ISR activity scores detectable at 0.1 mg/kg, a ∼50% reduction at 1 mg/kg, and near-complete normalization to WT levels at 10 mg/kg (**Fig. S4D-E**). DNL343 was also assessed in this study. A 1 mg/kg dose of RTX-117 more potently suppressed ISR pathway activation than a similar dose level of DNL343. The level of ISR pathway normalization achieved by 0.1 mg/kg RTX-117 approximated that of 1 mg/kg DNL343, consistent with the >10-fold potency advantage observed *in vitro* (**Fig. S4E** and **Fig. S2A**).

RTX-117 was well tolerated across all doses. Body weight, which is reduced in *Eif2b5* KI mice relative to WT animals, was partially preserved by RTX-117 treatment in a dose dependent manner (**Fig. S5A**). Motor coordination, assessed by beam walk, was significantly improved, with RTX-117-treated *Eif2b5* KI mice showing reduced traverse times and missteps relative to vehicle-treated *Eif2b5* KI mice (**Fig. S5B–C**). Histopathological analysis further confirmed RTX-117-driven attenuation of hallmark features of VWM disease neuropathology. Luxol Fast Blue staining revealed reduced myelin in vehicle-treated *Eif2b5* KI brain tissue relative to WT controls, consistent with previous reports (**Fig. S6A-B**). RTX-117-treated animals showed improvement in myelin integrity compared to vehicle controls (**Fig. S6A-B**), consistent with restoration of white matter architecture. GFAP immunoreactivity was elevated in *Eif2b5* KI mice and was nearly fully normalized by 1 and 10 mg/kg RTX-117 (**Fig. S6C-D)**. These data strongly indicate that pharmacologic activation of eIF2B by RTX-117 suppresses chronic ISR pathway activation in the brain and improves myelin integrity and motor function in a genetic model of ISR-driven neurodegeneration.

### eIF2B activator RTX-117 normalizes ISR pathway activation and improves neuromuscular function in a mouse model of CMT2D

Given the strong connection between ARS dysfunction, protein synthesis, and neurodegeneration, we next evaluated RTX-117 in *Gars*^P278KY/+^ mice, a model of CMT2D caused by a dominant pathogenic mutation in *GARS1*. *GARS1*, among the best-characterized ARS genes linked to CMT2 disease, is a nuclear-encoded glycyl–tRNA synthetase essential for protein synthesis in both the cytoplasm and mitochondria. Previous studies identified that protein synthesis is impaired in affected tissue of mice carrying *Gars1* mutations(*21, 23*), leading to GCN2 and ISR activation that drives neuropathy and motor dysfunction. Treatment began at weaning (3 weeks of age), when *Gars*^P278KY/+^ mice already show early motor impairment and axonal pathology. WT and *Gars*^P278KY/+^ mice were maintained on a RTX-117-or vehicle-formulated chow diet and were assessed longitudinally over eight weeks for behavioral and electrophysiological outcomes (**Fig. 3A**). A single oral dose (PO) of RTX-117 at 3 or 30 mg/kg in *Gars*^P278KY/+^ mice confirmed a favorable PK profile with expected dose-proportional exposure and kinetics (**Fig. S7A).** Under chow dosing at 30 and 300 mg/kg formulated chow (corresponding to estimated oral doses of 3 and 30 mg/kg), sustained systemic exposure of RTX-117 at both dose levels was confirmed by plasma PK throughout the study (**Fig. S7B**).

**Figure 3.**
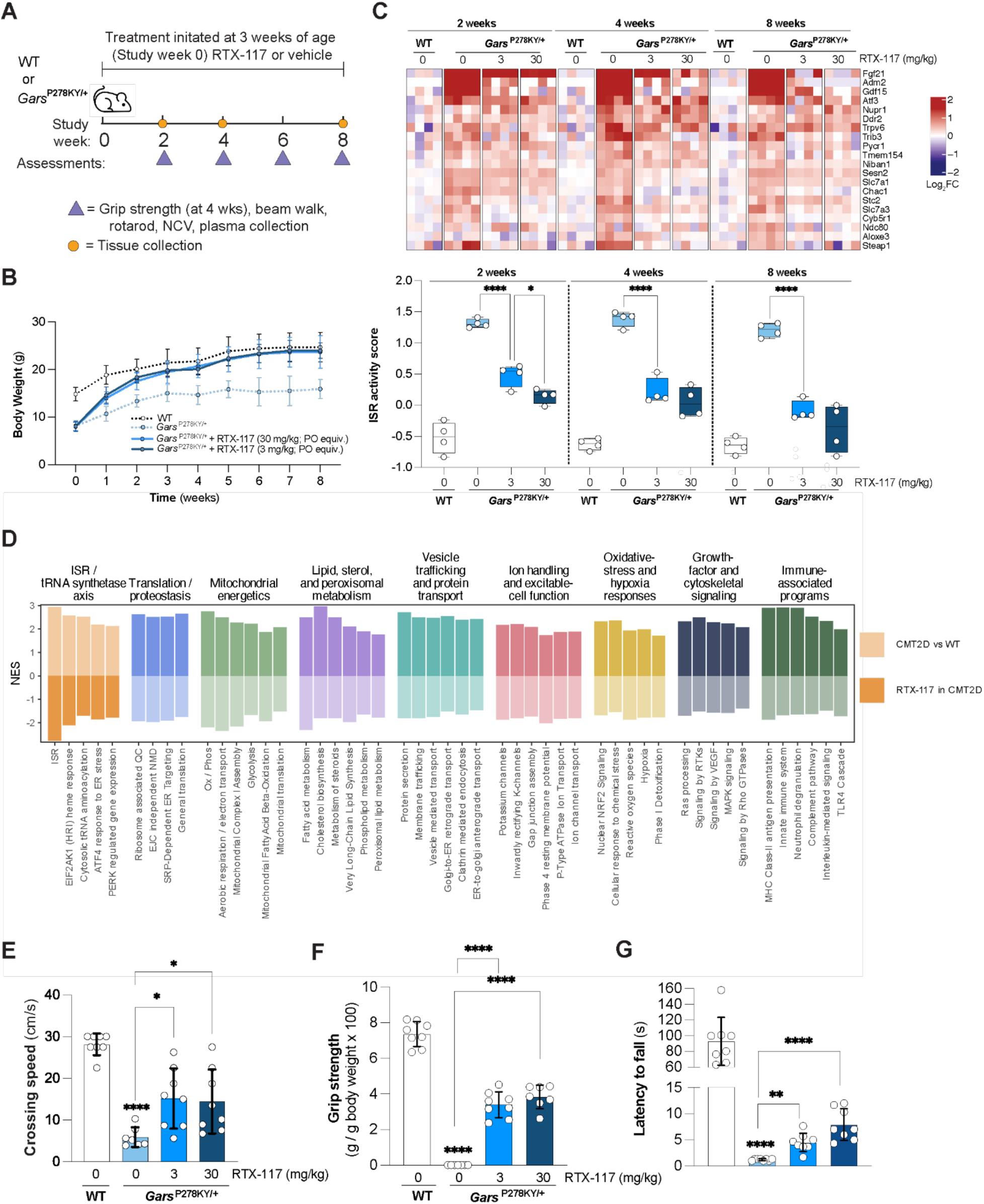
RTX-117 normalizes ISR pathway activation and improves neuromuscular function in a mouse model of CMT2D. (A) Experimental timeline and study design. Treatment with vehicle or RTX-117 formulated chow administration was initiated in *Gars*^P278KY/+^ mice at 3 weeks of age (study week 0). Functional assessments (beam walk, grip strength, rotarod, nerve conduction velocity) were performed longitudinally, and tissues were collected at the study endpoint. (B) Body weight trajectories of male and female wild type (WT) and *Gars*^P278KY/+^ mice during longitudinal treatment with RTX-117 or vehicle. (C) Heatmap of ISR pathway activation in spinal cord tissue from WT and *Gars*^P278KY/+^ mice treated with vehicle or RTX-117 at the indicated doses (mg/kg) across time points. Expression values are shown as log_2_ fold change relative to WT vehicle control from both male and female mice. The color scale represents log_2_ fold change, with red indicating increased and blue indicating decreased expression (top). ISR activity scores across treatment groups and time points, demonstrating normalization of ISR pathway activation with RTX-117 treatment. Points represent individual animals and boxes indicate median and interquartile range (bottom). (D) Pathways dysregulated in untreated *Gars*^P278KY/+^ mice relative to WT were identified by GSEA. Normalized enrichment scores (NES) are shown on the y-axis. Lighter colors denote the untreated *Gars*^P278KY/+^ vs WT comparison, whereas bolder colors denote the 30 mg/kg RTX-117 treated *Gars*^P278KY/+^ vs untreated *Gars*^P278KY/+^ comparison. For the pathways shown, the treatment-associated enrichment is in the opposite direction to the disease-associated enrichment, consistent with reversal toward the WT state. All displayed GSEA results met an FDR threshold < 10% (**Table S3G**). (E-G) Graphs showing crossing speed (beam walk), forelimb grip strength, and latency to fall (rotarod) at the 4-week post-treatment in WT, vehicle-treated *Gars*^P278KY/+^ mice, and RTX-117 treated *Gars*^P278KY/+^ mice. Data from males and females are presented as mean + SD. Statistical comparisons were performed using one-way ANOVA followed by Bonferroni’s multiple comparisons test. The *Gars*^P278KY/+^ mice vehicle groups were compared to WT vehicles and RTX-117 treatment groups. Brackets denote comparisons between *Gars*^P278KY/+^ groups. Significance for the WT versus *Gars*^P278KY/+^ mice vehicle comparison is indicated above the *Gars*^P278KY/+^ mice vehicle bar. *P < 0.05, **P < 0.01, ****P < 0.0001.

Vehicle-treated *Gars*^P278KY/+^ mice exhibited the expected lower body weight relative to WT littermates at 3 weeks of age and for the duration of the study (**Fig. 3B**). RTX-117 administration rescued body weight trajectories in a dose-dependent manner, with high-dose animals approaching WT body weight within two weeks of treatment and maintaining this gain for the duration of the study (**Fig. 3B**). To assess ISR pathway activation, we performed RNA-seq in spinal cord tissue from all groups. Although global differential expression profiles between CMT2D and WT mice were mild (**Fig S9A**), ISR activation was observed in *Gars*^P278KY/+^ mice relative to WT controls (**Fig S9B-G** and **Table S3E**), with upregulation of ATF4 target genes including *Fgf21*, *Gdf15*, *Chac1*, and *Trib3*, consistent with sustained translational stress in this model (**Fig. 3C, top**). Although *Gars1* expression has been previously associated with ISR activity (**Table S2**), no changes in *Gars1* mRNA levels were detected in CMT2D mice or with RTX-117 treatment (**Fig S9H**). As before, we constructed a 20-gene context-specific ISR signature and used it to quantify RTX-117 modulation in this study (**Table S3F** and methods). RTX-117 significantly suppressed ISR pathway activation in a dose-dependent manner, with reductions in ISR activity scores in *Gars*^P278KY/+^ mice detected at 2, 4, and 8 weeks post-treatment (**Fig. 3C, bottom,** and **Fig S9I**). After 8 weeks of treatment, high-dose RTX-117 normalized ISR activity scores to near WT levels, consistent with sustained normalization of translational stress signaling *in vivo* by treatment with an eIF2B activator. To our knowledge, these data are the first to demonstrate that pharmacological activation of eIF2B suppresses ISR pathway activation in a mouse model of CMT disease.

In addition to ISR dysregulation, a broader GSEA analysis using signatures from the Hallmark(*40*) and Reactome(*41*) databases identified 196 of 1,923 tested pathways as dysregulated pathways in *Gars*^P278KY/+^ mice compared with WT (FDR < 10%; **Table S3G**). Most (75%) of the dysregulated pathways were downregulated in *Gars*^P278KY/+^, which largely aligned with axes of biology related to tRNA aminoacylation, proteostasis, mitochondrial energetics, and lipid-related metabolism (**Fig 3D)**. Following RTX-117 treatment, the activity of 72% of the dysregulated pathways identified was reversed toward WT levels, suggesting that direct and specific reduction of the ISR pathway activation can improve other disease-associated phenotypes observed in *Gars*^P278KY/+^ mice (**Fig 3D)**.

Next, we assessed motor performance using beam-walk, grip strength, and rotarod tests. Consistent with prior reports, *Gars*^P278KY/+^ mice displayed severe impairment across all motor tests at baseline and throughout the study (**Fig 3E-G and Fig. S8**). RTX-117 treatment significantly improved coordination and strength as early as two weeks of treatment, in a dose-dependent manner, with the higher dose producing the more pronounced gains (**Fig. 3E-G**, **Fig. S8,** and **Movie S1-4**). *Gars*^P278KY/+^ mice continued to improve in motor performance and grip strength for the study duration (**Fig. 3E-G** and **Fig. S8**). In particular, grip strength showed the most substantial response, though rescue remained partial, likely reflecting the severity of the phenotype in *Gars*^P278KY/+^ mice at treatment initiation and the post-onset treatment start date. Earlier intervention or prolonged treatment may be required to achieve full rescue of motor dysfunction. These findings demonstrate that eIF2B activation by RTX-117 rescues core functional deficits in a mouse model of CMT2D, with the degree of improvement tracking with systemic drug exposure levels.

Finally, to assess whether functional improvements reflected recovery of peripheral axon integrity, we performed electrophysiological analyses of sciatic nerve conduction. *Gars*^P278KY/+^ mice exhibited the expected neuropathic deficits at baseline, including a 60% reduction in motor nerve conduction velocity (MNCV), reduced compound muscle action potential (CMAP) amplitude, and prolonged CMAP latency relative to WT mice (**Fig. 4**). After two weeks of RTX-117 treatment, MNCV improved modestly, reaching statistical significance in the high-dose group (**Fig. 4A-B)**. While distal CMAP amplitude was unchanged by RTX-117 treatment at two weeks, a directional trend towards improved latency was observed in the high dose group (**Fig. 4A and 4C-D)**. By 6 weeks, MNCV was significantly improved in a dose-dependent manner (**Fig. 4A and 4E)**, and CMAP latency was significantly decreased in RTX-117 treated *Gars^P278KY/+^*mice, with a trend toward improved CMAP amplitude at the high dose (**Fig. 4A and 4F-G)**. Proximal CMAP amplitude and latency showed a strong time- and dose-dependent improvement, with high-dose animals approaching WT conduction kinetics (**Fig. S10**). These findings demonstrate time- and dose-dependent recovery of motor axon function with RTX-117 treatment in *Gars*^P278KY/+^ mice, consistent with the dose-dependent ISR suppression, and support eIF2B activation as an effective *in vivo* modifier of ISR-driven peripheral neuropathy. Next, we assessed the levels of GDF15 and FGF21, two well characterized circulating factors and known targets of the ISR, in CSF and plasma of *Gars^P278KY/+^* mice after 8 weeks of RTX-117 treatment. Consistent with findings that GDF15 and FGF21 mRNA levels were significantly increased in *Gars^P278KY/+^* mice spinal cord tissue and reduced by RTX-117 treatment by RNA-seq, both GDF15 and FGF21 protein levels were increased in the CSF of *Gars*^P278KY/+^ mice relative to WT controls and were decreased in a dose-dependent manner by RTX-117 (**Fig. 4H-I**). No significant changes in plasma GDF15 and FGF21 were detected between WT and *Gars*^P278KY/+^ mice, with a trend towards decreased levels in the high dose treatment group (**Fig. S10F-G**). Of note, circulating GDF15 and FGF21 levels are reportedly elevated in CMT patients(*42–43*) suggesting that these ISR targets may serve as translational biomarkers for treatment response to RTX-117 in CMT patients.

**Figure 4.**
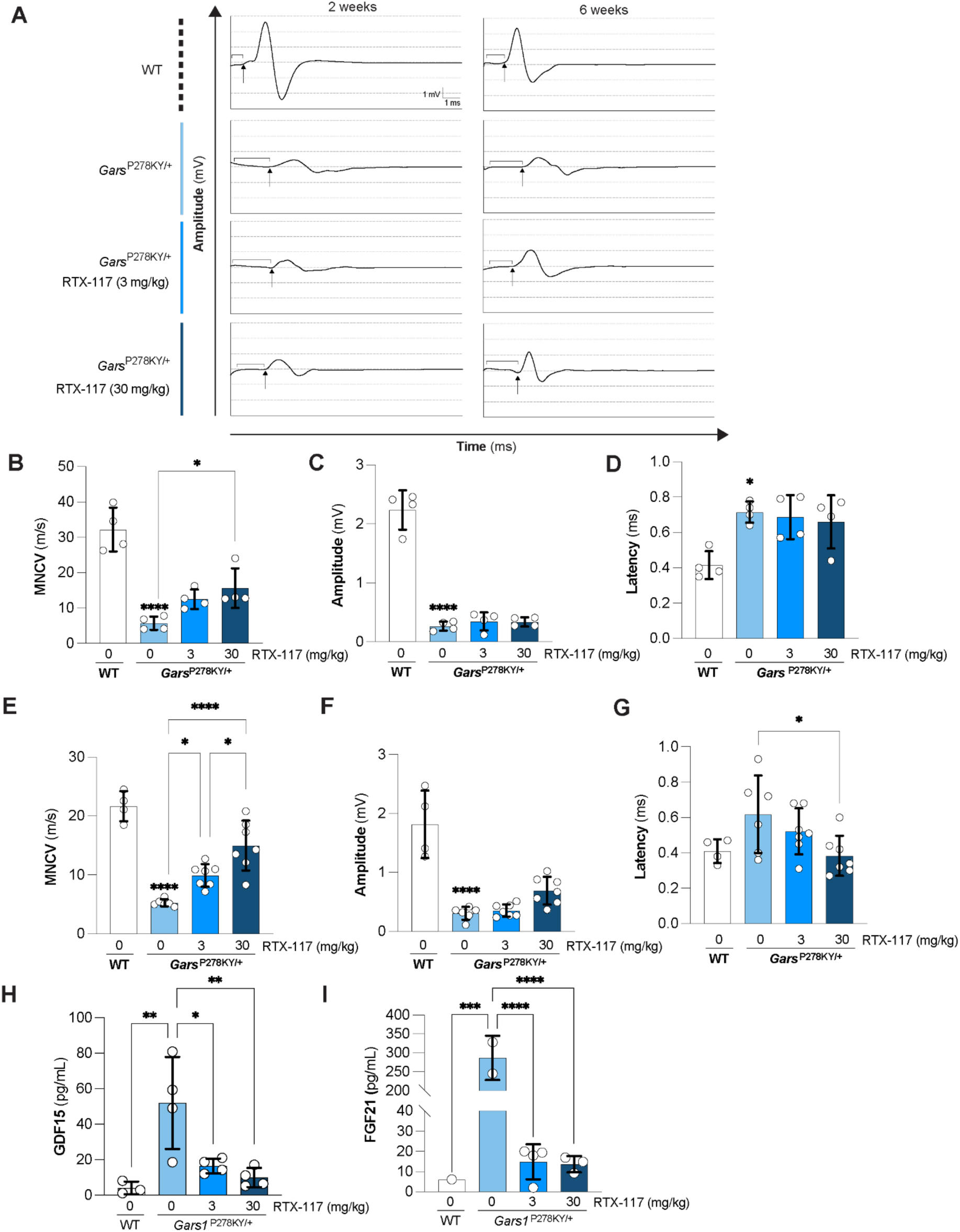
RTX-117 improves motor axon function and neuromuscular transmission in a mouse model of CMT2D. (A) Representative nerve stimulation evoked responses showing distal compound muscle action potential (CMAP) recording from wild type (WT) and *Gars*^P278KY/+^ mice at 2 (left) and 6 (right) weeks of treatment following stimulation at the achilles tendon. Scale bars, 1 mV, and 1 ms. Bracket denotes latency and arrow denotes onset. (B–D) Quantification of motor nerve conduction velocity (MNCV) (B), CMAP amplitude (C), and latency (D) in wild type (WT) and *Gars*^P278KY/+^ mice at 2 weeks post-treatment. (E–G) Quantification of (MNCV (E), CMAP amplitude (F), and latency (G) in wild type (WT) and *Gars*^P278KY/+^ mice at 6 weeks post-treatment. (H) GDF15 protein levels in CSF from wild type (WT) and *Gars*^P278KY/+^ mice following 8 weeks of RTX-117 treatment (I) FGF21 protein levels in CSF from wild type (WT) and *Gars*^P278KY/+^ mice following 8 weeks of RTX-117 treatment. Data from males and females are presented as mean ± SD. Statistical comparisons were performed using one-way ANOVA followed by Bonferroni’s multiple comparisons test. Untreated *Gars*^P278KY/+^ mice were compared to untreated WT mice and each RTX-117-treated group, and between the two RTX-117 dose groups. Brackets denote comparisons between *Gars*^P278KY/+^ groups. Significance for the WT versus *Gars*^P278KY/+^ comparison is indicated above the *Gars*^P278KY/+^ bar. *P < 0.05, **P < 0.01, ***P < 0.001, ****P < 0.0001.

### ISR pathway activation is a universal hallmark of tRNA synthetase dysfunction

Although ARS mutations account for the largest subset of CMT2 subtypes, it remains unclear whether chronic ISR activation is a unifying molecular hallmark across ARS-related neuropathies. To examine the prevalence of ISR activation beyond CMT2D, we assessed transcriptomic responses to loss-of-function perturbations against a broader range of cytoplasmic and mitochondrial ARS genes. To do this, we searched the NCBI Gene Expression Omnibus (GEO) for transcriptomic studies involving genetic or pharmacological loss of ARS function, including pathogenic mutations, knockout, knockdown, and enzymatic inhibition (**Table S4**). We then performed gene set enrichment analysis (GSEA)(*38–39*), comparing ARS-perturbed samples with their corresponding controls across multiple complementary ISR activation signatures (see “ISR Signature Collection” in methods section). Importantly, we noted that ISR activation in the spinal cord from *Gars*^P278KY/+^ mice in the present study was concordant with an independently generated transcriptomic dataset from the same mouse model(*23*), providing external validation of the data generated from this study (**Fig. 5A**, *GARS* ReviR vs sp. cord). Across the datasets examined, every ARS loss-of-function context showed significant enrichment of at least one ISR signature at an FDR threshold <= 10%, although the magnitude and breadth of activation varied across perturbation contexts (**Fig. 5A** and **Table S4).** Among the genetic perturbations, loss-of-function of *AARS2* in heart, *DARS2* in intestine, and *WARS1* in liver produced particularly strong ISR activation profiles. Additionally, we observed that pharmacological inhibition of tRNA synthetase activity with halofuginone or borrelidin induced robust ISR activation (**Fig. 5A**, last two conditions). Collectively, these findings indicate that ISR activation is a recurring consequence of impaired tRNA synthetase function across diverse genes, tissues, and perturbation modalities. Although ARS-associated forms of CMT are estimated to account for less than 2% of all CMT cases, this still represents more than 30,000-40,000 CMT patients worldwide (*4, 5, 44*). Beyond CMT, dominant or recessive variants in more than 30 ARS genes are associated with a broad spectrum of neurological, neuromuscular, and multisystem disorders, including spinal muscular atrophy, leukoencephalopathies, leukodystrophies, epilepsy, hereditary spastic paraplegia, myopathies, and mitochondrial disease phenotypes(*45, 46*).

**Figure 5.**
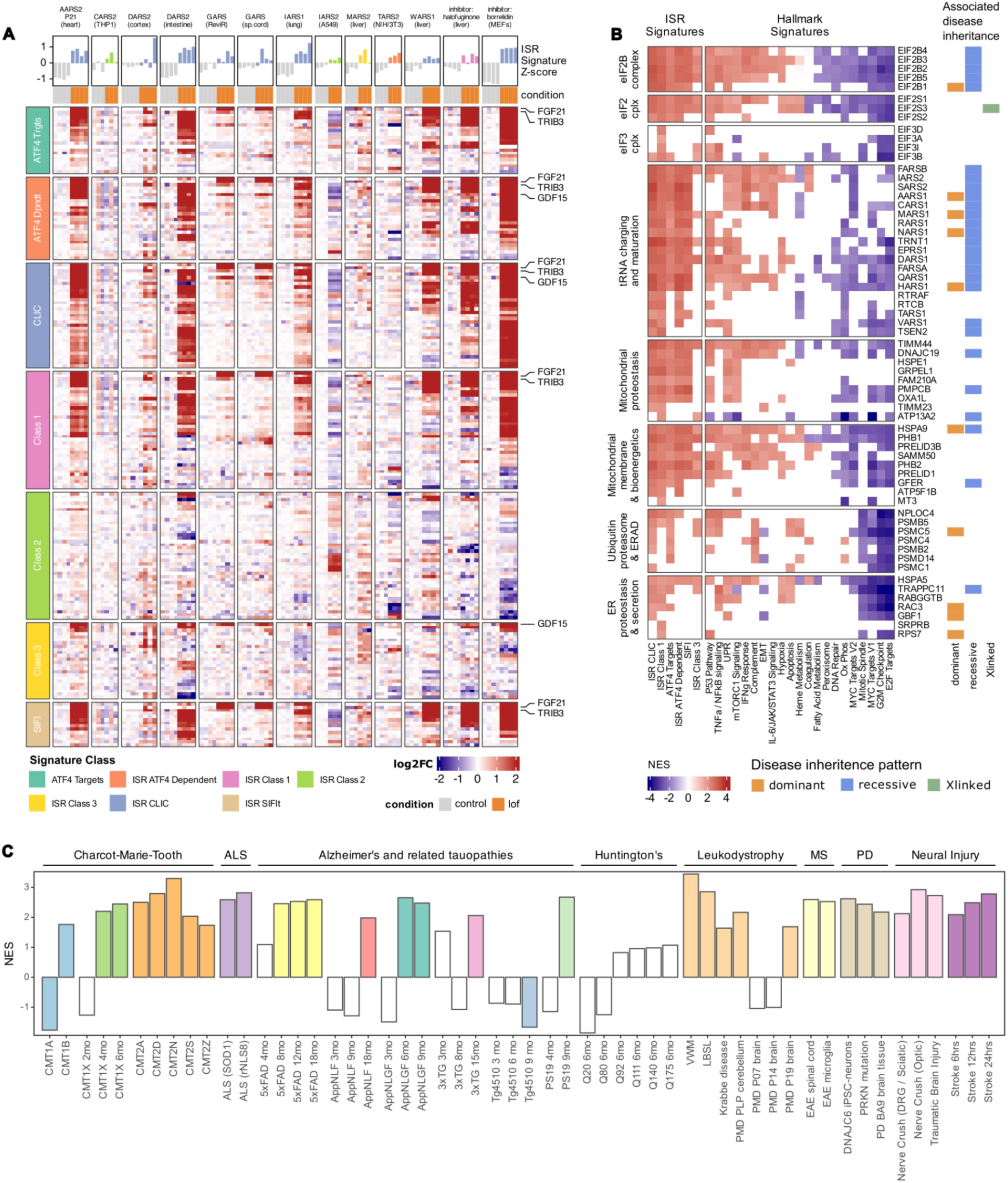
Uniform analytic framework identifies ISR pathway activation across genetic perturbations and disease models. (A) ISR activation following genetic or pharmacological disruption of aminoacyl-tRNA synthetase activity. The heatmap shows differential expression of genes from seven ISR signature sets. Columns represent individual samples grouped by NCBI GEO study, with control samples shown first and loss-of-function or pharmacologically perturbed samples shown second. Within each study, expression values were normalized to the mean of the corresponding control group and are displayed as log_2_ fold changes. Bars above the heatmap show z-transformed single-sample ISR signature scores and are colored according to the ISR signature class with strongest pathway enrichment. Each experiment displayed showed significant activation of the indicated ISR signature at FDR < 10%. Rows comprise the union of leading-edge genes identified by GSEA for each signature class across the experiments shown. (B) Pathway dysregulation inferred from Perturb-seq data. Tiles show the mean normalized enrichment score (NES) for each gene–pathway association across cellular contexts in which GSEA reached FDR < 10%. White tiles indicate that the pathway was not significantly altered in any analyzed cellular context. Genes are organized into functional groups derived from Gene Ontology enrichment analysis. GenCC annotations at right indicate genes with strong or definitive evidence of disease association, categorized by reported mode of inheritance: dominant (orange), recessive (blue), or X-linked (green). (C) GSEA analysis for ISR activity from different RNA-seq datasets, grouped by indication. Bar height indicates normalized enrichment score (NES) from GSEA analysis. Colored bars denote results that meet FDR < 10%. Abbreviations: ALS, amyotrophic lateral sclerosis; PD, parkinsonian disorders; MS, multiple sclerosis; LBSL, leukoencephalopathy with brain stem and spinal cord involvement and lactate elevation; PMD, Pelizaeus-Merzbacher disease.

### Perturb-seq associates ISR pathway-activating genes with monogenic-disease relevance

Having observed consistent ISR activation across manually identified datasets involving loss of aminoacyl-tRNA synthetase function, we next sought a more systematic approach to identify additional genes whose disruption may trigger ISR activation. We therefore analyzed publicly available Perturb-seq datasets from a variety of human cell lines curated through the EMBL-EBI Perturbation Catalogue(*47–50*). Within each experiment and cellular context, we assessed the effect of a gene knockout by performing GSEA using our ISR signature collection along with the MSigDB Hallmark gene signatures(*40*). Of 10,042 target genes represented in the source datasets, 389 genes had sufficiently broad transcriptional coverage amenable to our analysis approach (see methods and **Table S5A**). Among these were five genes associated with CMT: *AARS1* (CMT2N), *GBF1* (CMT2GG), *HARS1* (CMT2W), *MARS1* (CMT2U), and *NARS1* (CMT-NARS1)(*51*). Knockout of 102 of the 389 testable genes (26.2%), including all five CMT-associated genes mentioned above, produced positive enrichment of ISR signatures in at least one cellular context (**Fig. 5B** and **Fig. S11).**

Several functional classes of genes were associated with positive enrichment of ISR signatures upon knockout. For instance, knockout of each of the five subunits of the eIF2B complex was associated with ISR activation, as would be expected, as well as perturbations affecting components of the eIF2 and eIF3 complexes (**Fig. 5B** and **Table S5B**). Aminoacyl-tRNA synthetases and tRNA processing factors including *TRNT1*, *RTCB, and RTRAF* represented one of the largest functional classes, consistent with the results obtained from the bulk transcriptomic analysis (**Fig 5A)**. A second major class included genes involved in mitochondrial protein import, membrane organization, and respiration. Additional ISR-activating genes were associated with transcriptional regulation and protein homeostasis (**Fig. 5B** and **Fig. S11**). Analysis of Hallmark signatures revealed broader transcriptional programs accompanying these gene perturbations (**Table S5C**). Many ISR-activating perturbations were associated with enrichment of unfolded-protein-response, inflammatory, hypoxia, and apoptosis-related signatures (**Fig. 5B**). ISR activation was largely associated with reduction in cell growth and proliferation programs, as captured by negative normalized enrichment score (NES) from the MYC, E2F, G2M checkpoint, and mitotic-spindle-associated signatures. Mitochondrial perturbations additionally showed prominent negative enrichment of oxidative-phosphorylation signatures, consistent with the functional classification of these genes (**Fig 5B** and **Fig. S11)**. As several Hallmark gene sets share limited gene membership or correlation with the ISR signatures used in this analysis, these results describe transcriptional programs accompanying ISR activation rather than independent measurements of biochemical pathway activity.

To examine the disease relevance of the ISR-activating genes beyond CMT, we evaluated disease-gene associations curated by the *Gene Curation Coalition* (GenCC)(*52*). Among associations classified as having “strong” or “definitive” evidence, 41 of the 102 ISR-activating genes (40.2%) had at least one established disease association, compared with 74 of 287 genes (25.8%) not classified as ISR activating (**Table S5D**). ISR-activating genes were therefore significantly enriched for established monogenic or rare-disease associations (Yates-corrected χ² P=0.009; Fisher’s exact odds ratio =1.93, 95% CI 1.20–3.11). Together, these findings identify multiple forms of translation and stress pathways that converge on an ISR-activated transcriptional state and are triggered by loss-of-function mutations in genes often associated with rare and monogenic disease.

### ISR activation extends across genetically distinct CMT subtypes

Although the Perturb-seq analysis provided a systematic framework for identifying ISR-triggering loss-of-function perturbations, only five CMT-associated genes had sufficient transcriptional support for analysis. Notably, we could not assess perturbations to genes associated with some of the most prevalent forms of CMT, including CMT1A (*PMP22*), CMT2A (*MFN2*), and CMT1X (*GJB1*), an intermediate subtype with both axonal and demyelinating pathology(*53*). We therefore generated in-house RNA-seq data from mouse models of CMT2A (*Mfn2*^R94Q/+^), CMT1X (*Gjb1*-null), and CMT2N (AlaRS^R329H/+^) and searched NCBI GEO for additional CMT-relevant transcriptomic datasets. Inclusion of the CMT2N model allowed us to test whether ISR activation was reproducible across genetically distinct ARS-associated CMT subtypes.

GSEA of the in-house RNA-seq datasets identified significant ISR pathway activation in spinal cord from both CMT2N and CMT2A mice at two months of age (FDR <= 10%) (**Fig. 5C**). Longitudinal analysis of sciatic nerve from CMT1X mice at two, four, and six months revealed a progressive increase in ISR activation with age. Significant enrichment (FDR <= 10%) was first detected at four months and was more pronounced at six months, consistent with increasing ISR dysregulation during disease progression in this model. We additionally analyzed published RNA-seq datasets from models of CMT1A, CMT1B, CMT2S, and CMT2Z. Significant ISR pathway activation (FDR <= 10%) was detected in all models except CMT1A, where a reduction in ISR pathway activity was observed (**Fig 5C**, **Fig. S12** and **Table S6A**). Together, these results identify ISR activation as a recurring molecular feature across several genetically distinct axonal and intermediate CMT subtypes beyond those defined by ARS-associated dysfunction. The mixed ISR activity observed across demyelinating CMT models suggests that ISR modulation may also be relevant in select CMT1 subtypes, like CMT1B, but additional studies will be required to define the disease and cellular contexts in which it is therapeutically meaningful. Altogether our findings indicate a shared, ISR-associated molecular pathology across axonal CMT subtypes and support further exploration into normalization of ISR by eIF2B activation as a potential disease-modifying strategy.

### ISR activation is a recurrent feature across neurodegenerative diseases

Dysregulated eIF2α phosphorylation and ISR signaling have been reported across diverse neurodegenerative disorders. This evidence has accumulated through disease-specific studies using non-equivalent readouts, including PERK and eIF2α phosphorylation, induction of downstream effectors such as ATF4 protein levels, suppression of protein synthesis, and responses to genetic or pharmacological pathway modulation (*18, 19, 54*)(*55*). However, to our knowledge, a systematic analysis of the ISR pathway at the transcriptional level has not yet been reported across neurodegenerative disorders. We therefore sought to consolidate publicly available transcriptomic data from diverse neurological indications and determine whether ISR activation could be detected using a uniform analytical framework.

We manually curated human and mouse RNA-seq datasets from NCBI/GEO spanning amyotrophic lateral sclerosis (ALS), Alzheimer’s disease (AD) and related tauopathies, Huntington’s disease (HD), multiple sclerosis (MS), Parkinson’s disease and related parkinsonian disorders, leukodystrophies, and models of acute neural injury, including peripheral nerve crush, traumatic brain injury (TBI), and stroke. Within each study, samples were assigned to biologically relevant disease and control groups, and GSEA was performed using our set of primary and groomed ISR signatures (see methods, **Table S6-S7**). Significant ISR activation, defined at an FDR threshold <= 10%, was detected across most neurodegenerative disease categories and in many of the individual models examined, although substantial heterogeneity was observed (**Fig. 5C**). By applying the same ISR signatures, enrichment procedure, and significance criterion across independently generated studies, this analysis provides a consolidated view of ISR transcriptional activity across neurological conditions that have previously been investigated largely in isolation (**Fig. 5C**).

As expected, ISR activation was detected in rare inherited leukodystrophies including VWM, leukoencephalopathy with brainstem and spinal cord involvement and lactate elevation (LBSL), Pelizaeus-Merzbacher Disease (PMD), as well as in mouse models of ALS (**Fig. 5C**). Significant ISR activation was also detected in postmortem brain samples from individuals with Parkinson’s disease compared with neurologically healthy controls (**Fig. 5C**, PD BA9 brain tissue). In contrast, we did not detect significant ISR enrichment in expression profiles from the striatum of HD mouse models with increasing CAG repeat lengths, despite reports of increased PERK-dependent eIF2α phosphorylation in HD models(*56*). A prominent temporal pattern of ISR activation was observed across multiple mouse models of AD and related tauopathies. This was most clearly observed in 5xFAD mice, where ISR activation was first detected at eight months and increased in magnitude at 12 and 18 months where advanced cognitive decline and disease pathology is observed. In other amyloid-predominant models, such as App^NL-F^, and App^NL-G-F^, significant ISR enrichment was also detected by the last time point assayed. Similar patterns were identified in the 3xTg-AD model, which develops both amyloid and tau pathology, as well as the PS19 tauopathy model. In the Tg4510 tauopathy model, however, the pattern of ISR activity was reversed at the endpoint. Thus, our uniform analysis across these independent datasets revealed an age-associated emergence of ISR activity spanning amyloid-predominant, combined amyloid–tau, and tau-predominant pathologies (**Fig. 5C**).

## DISCUSSION

Herein, we demonstrate that ISR pathway activation drives axonal dysfunction in a mouse model of CMT2D and that pharmacological eIF2B activation with RTX-117 treatment normalizes translational homeostasis, rescues motor performance, and improves nerve conduction after disease onset. Beyond CMT2D, we establish ISR pathway activation as a molecular hallmark of tRNA synthetase dysfunction and a shared feature of additional CMT subtypes associated with, for example, mitochondria and myelin dysfunction. These findings support a model in which maladaptive translational repression is a functional driver of axonal pathology in CMT subtypes rather than merely a marker of cellular stress. More broadly, our results pinpoint the eIF2ɑ-ATF4 axis as a therapeutically tractable target linking translational control to axonal degeneration. Normalizing chronic ISR activation through eIF2B stabilization may therefore represent a disease-modifying strategy for axonopathies in which translational stress is a central pathogenic mechanism. GDF15 and FGF21, circulating ISR target genes are emerging as diagnostic biomarkers for multiple CMT subtypes including CMT2D(*42*)(*23*), are modulated by RTX-117 treatment in CMT2D mice, and may serve as a valuable pharmacodynamic markers for RTX-117 response across genetically heterogeneous neuropathies. Notably, the genetic driver remains unknown in approximately >20% of CMT cases, a population that may also harbor uncharacterized ISR-activating mechanisms.

tRNAs play an essential role in protein synthesis, with disruption in mRNA translation increasingly recognized as a driver of human diseases associated with tRNA dysfunction(*57*). Mutations in cytoplasmic and mitochondrial tRNA synthetases underlie a spectrum of neurodegenerative diseases including CMT, ataxia, leukoencephalopathies, and leukodystrophies. Our data establish ISR pathway activation as a shared molecular consequence of tRNA synthetase dysfunction, consistent with prior work linking ARS mutations to translational stress. The particular vulnerability of the nervous system to ARS dysfunction may reflect the elevated proteostasis demands of long-lived, metabolically active neural cell types within the CNS and periphery. While tRNA synthetase mutations are believed to cause disease through partial loss of aminoacylation activity and, in some cases, gain-of-toxic-function mechanisms such as pathological aggregation or aberrant protein-protein interactions, it is clear that eIF2ɑ-ATF4 axis activation is a downstream consequence of proteotoxic stress that drives neurodegeneration. Mutations in GARS1 activate the ISR through the stress sensing kinase GCN2(*23*), and GARS1 dysfunction induces ribosome pausing at glycine codons reliant on tRNA^Gly^ for decoding(*21*). It is likely that ARS-associated mutations in other CMT2 subtypes also activate GCN2 and the ISR pathway by a similar mechanism. Indeed, deletion of GCN2 inhibits ISR pathway activation and alleviates neuropathy in a mouse model of CMT2D(*23*). Our data support this model and demonstrate that pharmacological restoration of eIF2B activity normalizes downstream stress signaling and improves axonal physiology. For ARS genes with mitochondrial functions, including GARS1, KARS1, and IARS2, ISR activation may additionally arise through the DELE1- HRI axis in response to mitochondrial stress(*58*). Another possibility is that perturbations in mitochondrial proteostasis activate the mitochondrial unfolded protein response (UPR^mt^) pathway that may further contribute to translational suppression and metabolic dysfunction(*59*), a hypothesis further supported by our reanalysis of Perturb-seq data that showed that perturbations to mitochondrial genes comprised one of the largest functional categories of genes that trigger ISR activation. It is worth noting that mutations in several factors involved in mitochondrial function and translation, intracellular trafficking, and vesicle transport also cause axonal CMT and may engage overlapping stress response mechanisms.

A key question remains as to why specific disease states or neural cell types such as sensory or motor neurons show particular vulnerability to ISR pathway activation. One possibility is that the maintenance of axonal integrity requires continuous synthesis and renewal of proteins involved in cytoskeletal maintenance, axonal transport, and mitochondrial function. Indeed, several studies implicate mRNA transport from the cell body to distal axonal compartments to support eIF2-dependent localized mRNA translation particularly upon stress and at neuromuscular junction and includes synthesis of proteins such as Gap43, Stat3, Agrin, Nmnat2, and additional factors associated with glucose metabolism, synaptic connection, and protein homeostasis(*60–62*). Given the extraordinary length and metabolic demands of peripheral axons, chronic translational repression likely imposes a disproportionate burden on distal axonal homeostasis. Restoring eIF2B complex activity may restore translational capacity while preserving upstream stress sensing, providing a mechanism to rebalance protein synthesis without ablating the adaptive stress response. Whether ISR pathway activation contributes to pathology across other axonal neuropathies remains to be determined, but our data support ISR normalization as a broad therapeutic strategy in axonopathies where translational stress is a central component of disease pathogenesis.

Our unbiased profiling of disease associated transcriptomics datasets identify a number of disorders with elevated ISR pathway activation including VWM, CMT, and ALS. Clinical-stage compounds that modulate eIF2B, such as DNL343 and fosigotifator, have demonstrated central nervous system penetration and appeared well tolerated in early clinical studies(*34*)(*35*), though whether eIF2B activation confers clinical benefit remains to be established in ongoing VWM clinical trials. Studies in ALS patients highlights the complexity of translating ISR modulation into efficacy. Furthermore, despite clear ISR pathway activation in select ALS subtypes such as TDP43 and SOD1 mutant preclinical models, efficacy studies following eIF2B activation are lacking. Together, these findings indicate that the degree of ISR pathway dependency may vary across disease contexts and warrants evaluation in disease-relevant models. Developed to overcome the PK limitations of previous eIF2B activators, RTX-117 provides enhanced potency and favorable drug-like attributes required to maintain translational homeostasis in patients with neurological disorders. In biochemical and cellular assays, RTX-117 demonstrates substantially increased potency relative to first-generation clinical eIF2B activators and robust suppression of ISR pathway activation *in vivo*. The efficacy observed in the CMT2D mouse model described herein may therefore reflect both the proximal relationship between ISR activation and disease pathogenesis in this CMT subtype and the ability of RTX-117 to durably restore eIF2B function and attenuate established neuropathy. These findings raise the possibility that disease context and the degree of ISR dependency are important determinants of therapeutic responsiveness to eIF2ɑ-ATF4 axis targeted therapies.

Beyond peripheral neuropathy, accumulation of misfolded amyloid-β and tau proteins in Alzheimer’s disease triggers ISR signaling through upstream kinases, leading to phosphorylation of eIF2α, suppressing global protein synthesis, and driving translational reprogramming(*30, 63–65*). Consistent with these findings, our analysis provides strong evidence of chronic ISR pathway activation across a number of genetically distinct mouse models of AD. Chronic ISR activation in these contexts may impair synaptic plasticity and neuronal maintenance processes that depend on sustained protein synthesis. Nevertheless, the presence of ISR activation across neurodegenerative disorders does not necessarily imply uniform pathway dependency and further preclinical and clinical evaluation are warranted. Overall, our findings establish maladaptive translational repression as a modifiable driver of axonal dysfunction in ISR-proximal neuropathies and provide a strong mechanistic rationale for the clinical evaluation of RTX-117 in genetically defined CMT2 patient populations characterized by ISR activation.

## MATERIALS AND METHODS

### Compound design and chemical synthesis

DNL343 was synthesized as previously described(*66*). The synthetic details for preparation of RTX-117 have been reported previously (US patent 12,612,392). The synthetic route is summarized in **Figure S13,** and full procedures are described below.

### Step A: methyl 2-(4-chloro-3-fluorophenoxy)acetate (2)

To a solution of 4-chloro-3-fluorophenol (25.0 g, 171 mmol) in DMF (100 mL) was added methyl bromoacetate (31.3 g, 205 mmol) and potassium carbonate (71.7 mg, 511 mmol) under N_2_ for 16 hours. The reaction mixture was poured into water, 80°r (200 mL) and extracted with EtOAc (200 mL x 3). The combined organic phase was washed with brine (50 mL), dried over anhydrous Na_2_SO_4_, filtered and concentrated to dryness under reduced pressure to give the residue. The residue was purified by silica gel chromatography (eluent: petroleum ether: EtOAc = 5: 1 to 10 : 1) to afford methyl 2-(4-chloro-3-fluorophenoxy)acetate (33 g, 89.18 %) as yellow oil. Mass Spectrum (ESI) m/z = 219.0 (M+H) +.

### Step B: 2-(4-chloro-3-fluorophenoxy)acetic acid (3)

To a solution of methyl 2-(4-chloro-3-fluorophenoxy)acetate (20.0 g, 91.5 mmol) in tetrahydrofuran /H_2_O(100 mL, V= 5:2) was added lithium hydroxide (4.38 g, 182.9 mmol) and stirred for 18 hours at 25 °C. Adjust the pH with 2 M HCl to ∼3, extracted with EtOAc (30 mL x3). The combined organic phases were washed with brine (50 mL x2), dried over anhydrous Na_2_SO_4_, filtered and concentrated to dryness under reduced pressure to give the residue. The residue was purified by silica gel chromatography (eluent: ACN: water(1‰ FA) = 5: 1 to 20 : 1) to afford 2-(4-chloro-3-fluorophenoxy)acetic acid as white solid (18 g, 86.55%). Mass Spectrum (ESI) m/z = 203.0(M-H) ^-^

### Step C: methyl (1R,4R)-4-(2-(4-chloro-3-fluorophenoxy)acetamido)cyclohexane-1-carboxylate (4)

To a solution of 2-(4-chloro-3-fluorophenoxy)acetic acid (26.0 g, 127.2 mmol)in DMF (20 mL) was added HATU (72.5 g, 190.8 mmol), DIEA (49.2 g, 381.6 mmol), methyl (1R,4R)-4-aminocyclohexane-1-carboxylate (20.0 g, 127.2 mmol), the reaction was at ambient temperature and stirred for 16 hours. The reaction mixture was poured into water (200 mL) and extracted with EtOAc (200 mL x 3). The combined organic phase was washed with brine (50 mL), dried over anhydrous Na_2_SO_4_, filtered and concentrated to dryness under reduced pressure to give the residue. The residue was purified by silica gel chromatography (eluent: petroleum ether: EtOAc = 5: 1 to 10: 1) to give methyl (1R,4R)-4-(2-(4-chloro-3-fluorophenoxy)acetamido)cyclohexane-1-carboxylate (36 g, 74.08 %) as yellow oil. Mass Spectrum (ESI) m/z = 344.0 (M+H) ^+^.

### Step D: 2-(4-chloro-3-fluorophenoxy)-N-((1R,4R)-4-(hydrazinecarbonyl)cyclohexyl) acetamide (5)

To a solution of methyl (1R,4R)-4-(2-(4-chloro-3-fluorophenoxy)acetamido)cyclohexane-1-carboxylate (20 g, 58.1 mmol) in ethanol (100 mL) was added N_2_H_4_.H_2_O (29.0 g, 290.8 mmol), the reaction was at 80 °C and stirred 18 hr. The reaction mixture concentrated to dryness under reduced pressure to give the residue. The residue was recrystallized with ACN to afforded 2-(4-chloro-3-fluorophenoxy)-N-((1R,4R)-4-(hydrazinecarbonyl)cyclohexyl) acetamide (19 g, 95 %) as white solid. Mass Spectrum (ESI) m/z = 344.2(M+H) ^+^.

### Step E: 2-(4-chloro-3-fluorophenoxy)-N-((1R,4R)-4-(5-hydroxy-1,3,4-oxadiazol-2-yl) cyclohexyl)acetamide (6)

To a solution of 2-(4-chloro-3-fluorophenoxy)-N-((1r,4r)-4-(hydrazinecarbonyl)cyclohexyl) acetamide (20 g, 58.1 mmol) in DCM (100 mL) was added CDI (28.2 g, 174.5 mmol), the reaction was at ambient temperature and stirred 16 hr. The reaction mixture was poured into water (200 mL) and extracted with EtOAc (200 mL x 3). The combined organic phase was washed with brine (50 mL), dried over anhydrous Na_2_SO_4_, filtered and concentrated to dryness under reduced pressure to give the residue. The residue was purified by silica gel chromatography (eluent: petroleum ether: EtOAc = 3: 1 to 10: 1) to afford 2-(4-chloro-3-fluorophenoxy)-N-((1R,4R)-4-(5-hydroxy-1,3,4-oxadiazol-2-yl)cyclohexyl)acetamide (20.1 g, 84.09 %) as white solid. Mass Spectrum (ESI) m/z = 370.0 (M+H) +.

### Step F: benzyl 3-(trifluoromethoxy)azetidine-1-carboxylate (8)

To a solution of benzyl 3-hydroxyazetidine-1-carboxylate (20 g, 96.5 mmol) in EtOAc (500 mL) was added AgOTf (74.41 g, 289.5 mmol), Selectflour (6.4 g, 18.1 mmol), potassium fluoride (22.4 g, 386.0 mmol), 2-fluoropyridine (26.9 g, 289.5 mmol), TMSCF3 (41.1 g, 289.5 mmol), the reaction was at ambient temperature and stirred 18 hr. TLC show the reaction was completed. The reaction mixture was poured into water (100 mL) and extracted with EtOAc (100 mL x 3). The combined organic phase was washed with brine (50 mL), dried over anhydrous Na_2_SO_4_, filtered and concentrated to dryness under reduced pressure to give the residue. The residue was purified by silica gel chromatography (eluent: petroleum ether: EtOAc = 3: 1 to 10: 1) to afford benzyl 3-(trifluoromethoxy)azetidine-1-carboxylate(10.0 g, 37.6%) as yellow oil. 1H NMR (400 MHz, CDCl_3_) δ 7.28 –7.17 (m, 5H), 5.02 (s, 2H), 4.84 – 4.81 (m, 1H), 4.24 – 4.19 (m, 2H), 4.06 – 4.02 (m, 2H).

### Step G: 3-(trifluoromethoxy)azetidine.HCl (9)

To a solution of benzyl 3-[(trifluoromethyl)oxy]azetidine-1-carboxylate (10.0 g, 36.3 mmol) in MeOH (50 mL) was added Pd/C (1.93 g, 18.1 mmol), con. HCl (8 mL, 32.0 mmol), the reaction was at 50 °C and stirred 16 hr under an atmosphere of H_2_ . The mixture was filtered and concentrated to dryness under reduced pressure to give the crude product 3- (trifluoromethoxy)azetidine.HCl (2.96 g, 46%) as a colorless oil.

### Step H: RTX-117 2-(4-chloro-3-fluorophenoxy)-N-((1R,4R)-4-(5-(3- (trifluoromethoxy)azetidin-1-yl)-1,3,4-oxadiazol-2-yl)cyclohexyl)acetamide

To a solution of 2-(4-chloro-3-fluorophenoxy)-N-((1R,4R)-4-(5-hydroxy-1,3,4-oxadiazol-2- yl)cyclohexyl)acetamide (11 g, 29.7 mmol) in DMF (30 mL) was added 3-(trifluoromethoxy)azetidine.HCl (5.0 g, 35.6 mmol), BOP (19.6 g, 44.6 mmol), DIEA (15.3 g, 118.8 mmol), the reaction was at ambient temperature and stirred 16 hr. The mixture was poured into water (100 mL) and extracted with EtOAc (100 mL x3). The combined organic phase were washed with brine (50 mL x2), dried over anhydrous Na_2_SO_4_, filtered and concentrated to dryness under reduced pressure to give the crude product. The crude produce was subjected to Prep-HPLC ((ACN : water (1‰ FA)) to afford 2-(4-chloro-3-fluorophenoxy)-N-((1R,4R)-4-(5- (3-(trifluoromethoxy)azetidin-1-yl)-1,3,4-oxadiazol-2-yl)cyclohexyl)acetamide (13 g, 89 %) was obtained as a white solid.

LCMS: m/z =493.2 (M+H)+.

1H NMR (400 MHz, DMSO) δ 8.01 (d, J = 8.0 Hz, 1H), 7.50 (t, J = 8.9 Hz, 1H), 7.07 (dd, J = 11.4, 2.8 Hz, 1H), 6.91 – 6.74 (m, 1H), 5.32 – 5.31 (m, 1H), 4.51 (s, 2H), 4.47 –4.43 (m, 2H), 4.20 – 4.17 (m, 2H), 3.83 – 3.57 (m, 1H), 2.76 – 2.67 (m, 1H), 2.04 – 2.01 (m, 2H), 1.92 – 1.81 (m, 2H), 1.63 – 1.45 (m, 2H), 1.41 – 1.09 (m, 2H).

### Cryo-EM structure determination of RTX-117 and visualization

Structural work was provided as a service by Biortus Biosciences Co. Ltd (Wuxi, China). Genes encoding the five subunits of human eIF2B were synthesized, cloned into the pcDNA3.4 vector, and sequenced to confirm sequences. The α subunit plasmid was transfected individually into HEK293F cells, while the plasmids encoding the remaining four subunits (β, γ, δ, ε) were co-transfected at a molar ratio of 1:1:1:1. Cells were cultured at 37°C with 5% CO₂ for 72 hours and following incubation, cells were harvested and resuspended in 220 mL lysis buffer (20 mM HEPES-KOH pH 7.5, 200 mM KCl, 1 mM TCEP, 5 mM MgCl₂, nuclease, and complete Protease Inhibitor Cocktail, Roche). The suspension was mixed at 4°C and lysed by high-pressure homogenization (microfluidics homogenizer, 650 bar, 4 passes). Lysate was centrifuged at 16,000 rpm for 60 min at 4°C, and the supernatant was collected for affinity chromatography. The target proteins were captured using a Ni Bestarose FF column and eluted with a step gradient of 10, 20, 100, and 300 mM imidazole. The 100 mM and 300 mM fractions were combined and dialyzed into Buffer A (20 mM HEPES-KOH pH 7.5, 50 mM KCl, 1 mM TCEP, and 5 mM MgCl₂). The dialyzed sample was loaded onto an 8 mL Capto HiRes Q 10/100 GL column (GE Healthcare), washed with Buffer A, and eluted over a 400 mL linear gradient of 50-600 mM KCl. The eIF2B fractions eluted at ∼410 mM KCl. Fractions were pooled and concentrated using an Amicon Ultra- 15 centrifugal filter (50 kDa MWCO). Concentrated fractions were loaded onto a 24 mL Superdex 200 10/300 GL column (GE Healthcare) pre-equilibrated with Buffer C (20 mM HEPES-KOH pH 7.5, 100 mM KCl, 1 mM TCEP, and 5 mM MgCl₂). The α subunit and the other four subunits were mixed at a molar ratio of 1.5:1 and incubated on ice for 1.5 h to assemble the decameric complex. The assembled complex was then purified via a 24 mL Superose 6 Increase 10/300 GL column pre-equilibrated with Buffer C. The final yield was ∼1 mg per liter of cell culture.

Au 300-mesh R1.2/1.3 holey carbon grids (Quantifoil) were glow-discharged for 100 s at 15 mA (PELCO easiGlow). The eIF2B complex was centrifuged at 12,000 × g for 15 min before grid preparation. A 3.5 µL aliquot (1.5 mg/mL) was applied to each grid, blotted for 3.5s at a blot force of 1 at 4°C under 100% humidity and immediately plunge-frozen in liquid ethane using a Vitrobot Mark IV (Thermo Fisher Scientific). Cryo-EM data were collected on a 300 kV Titan Krios G4 electron microscope (Thermo Fisher Scientific) equipped with a K3 direct electron detector and a Selectris X Imaging Filter (20 eV slit). Data collection was automated using EPU software. Movies were recorded in super-resolution mode (2x binning, physical pixel size 0.819 Å) at a nominal magnification of 105,000×. Total electron dose was 56 e⁻/Å² distributed across 50 frames. A total of 2,030 movies across 2 batches (530 and 1500) with a defocus range of -1.1 µm to -2.4 µm. All image processing was performed using cryoSPARC v4.6.0 unless otherwise noted. After patch motion correction and patch CTF estimation, 476,983 particles (86,424 + 390,559) were picked by template-based picking and extracted with 2×2 binning (1.638 Å/pixel, 226-pixel box size). Two rounds of 2D classification isolated high-quality particles, which were used for ab initio reconstruction. Heterogenous refinement into three classes identified a dominant well-resolved class (Class 2), which was selected for homogeneous refinement. Refined coordinates were used to re-center and re-extract 263,354 unbinned particles (0.819 Å/pixel, 452-pixel box size). These particles underwent non-uniform refinement followed by local refinement, yielding a final map at 2.82 Å resolution. Structural data was analyzed using The PyMOL Molecular Graphics System, Version 3.1 Schrödinger, LLC.

### Fluorescence Polarization Assay

The fluorescence polarization (FP) assay was performed in reaction buffer containing 20 mM HEPES (pH 7.5), 100 mM KCl, 5 mM MgCl₂, 1 mM TCEP, 0.1 μM FAM-labeled RTX-117, 2% DMSO, and 100 nM purified eIF2B complex (described above). Unlabeled RTX-117 were prepared as a 0.2 mM DMSO stock and serially diluted 3-fold using an Echo acoustic liquid handler to generate 10-point concentration-response curves (top final concentration: 500 nM). RTX-117 was dispensed into 384-assay plates, followed by addition of FAM-RTX-117 working solution and the eIF2B complex. Reactions were incubated at room temperature for 60 min. Fluorescence polarization was measured using a TECAN M1000 microplate reader with the following setting; excitation: 470 nm, emission: 530 nm, G-factor: 1.089.

### Cell culture

HEK293T cells (ATCC, CRL-3216) were cultured in DMEM (Corning) supplemented with 10% FBS and 1% penicillin-streptomycin. A HEK293T cell line stably expressing an ATF4 uORF- luciferase reporter was generated as previously described(*28*). ATF4-luciferase reporter HEK293T cells were seeded into 96-well plates (Corning, 3916) in 100 µL media. After overnight incubation, cells were co-treated with 50 nM thapsigargin (MCE, HY-13433) and RTX-117 at varying concentrations and incubated at 37°C for 6 hours. DMSO-treated cells served as the vehicle control and cells treated with 50 nM thapsigargin alone served as the positive control. Plates were equilibrated to room temperature for 10 min before measuring luciferase activity with the ONE-Glo Luciferase Assay System (Promega) per the manufacturer’s instructions. Luminescence was detected on a Varioskan LUX Microplate Reader (Thermo Fisher Scientific). Normalized luciferase activity was calculated using the following formula:

Normalized Activity =Read - NC PC - NC 100

IC50 values were determined using non-linear regression analysis in GraphPad Prism 10.

For ATF4 protein western blot analysis, HEK293T cells were seeded into 12-well plates and incubated overnight at 37°C with 5% CO₂. The cells were co-treated with 50 nM thapsigargin and varying concentrations of RTX-117 for 2 hours.

### Immunoblot analysis of ATF4

Cells were washed with PBS and lysed in RIPA Buffer. Lysates were denatured at 95°C for 15 min, resolved by SDS-PAGE, and transferred onto PVDF membranes. Membranes were incubated overnight at 4°C with primary antibodies against ATF4 (Cell Signaling Technology, 11815; 1:1000 dilution) and GAPDH (Proteintech, 60004-1-Ig; 1:1000). After washing with TBST, membranes were incubated with HRP-conjugated secondary antibodies (Cell Signaling Technology, 7074P2 and 7076P2) for 1 hour at room temperature. Signals were detected with the BeyoECL Moon kit (Beyotime, P0018FM) on a Tanon T1600 imager. Band densitometry was performed in ImageJ software.

### ISR induction in primary human PBMCs

PBMCs from four healthy donors were obtained from Meisen CTCC (Cat. # WEZ4148, WEZ4021-1, W013K144, WEZ4125). PBMCs were thawed and recovered in HBSS (ATCC, 30- 2213) supplemented with 10% FBS (EXcell, FSP500) and seeded into 6-well plates at 2x10^6^ cells per well in 2 mL. After overnight incubation at 37°C with 5% CO₂, cells were treated with RTX- 117 at the indicated conditions (0.1% DMSO final) for 22 hours to allow compound engagement before stress induction. Thapsigargin was then added to 50 nM and incubated for 2 hours. PBMCs were harvested by centrifugation at 200 × g for 5 min, and total RNA was extracted using the RNeasy Plus Mini Kit (Qiagen, 74136) per the manufacturer’s instructions.

### Animal care

Animals were housed under a standard 12-hour light/dark cycle with ad libitum access to food and water. All animal procedures were conducted and approved by WuXi AppTec (Nantong, China) or XtalPi Inc. Institutional Animal Care and Use Committee or Ethics Committee (IACUC). During the study, the care and use of animals were conducted in accordance with the regulations of the Association for Assessment and Accreditation of Laboratory Animal Care (AAALAC).

### VWM Mouse Model

The VWM knock-in mouse line carrying a p.R191H substitution in the *Eif2b5* gene (Ensembl: ENSMUSG00000003235) was generated using CRISPR/Cas9-mediated homologous recombination as a service at Shanghai Model Organisms. Briefly, Cas9 mRNA, guide RNA (targeting sequence: 5’-ACGCTGCCATGAGGACAACG-3’), and a synthetic oligo donor DNA carrying the target point mutation (along with synonymous mutations to prevent secondary Cas9 cleavage) were co-microinjected into the fertilized eggs of C57BL/6J mice. The resulting F0 founder mice were crossed with wild-type C57BL/6J mice to generate F1 heterozygous offspring. Heterozygous mice were then intercrossed to obtain homozygous knock-in animals and wild-type (WT) littermate controls for all experiments. Animals were genotyped using genomic DNA extracted from tail clips by PCR amplification using specific primers (Forward: 5’- CAGAGCCCTGGAGGAACAC-3’; Reverse: 5’-TGGAACAGGCTCTGAAGGGA-3’), followed by direct Sanger sequencing of the PCR products to confirm the presence of the mutation.

### CMT2D mouse model

CMT2D animal work was provided as a service by WuXi AppTec (Shanghai, China). *Gars^P278KY/+^* mice were obtained from the Jackson Laboratory (B6; CAST-Gars^Nmf249^/RwbJ, JAX stock #033165). IVF was performed for colony expansion using 1 male heterozygous *Gars^P278KY/+^* mouse and 4-week-old female C57BL/6 mice (Charles River Laboratories). Genotypes were confirmed by PCR analysis of genomic DNA isolated from tail clips as previously described. *Gars^P278KY/+^* mice were divided into 3 groups based on their weights after finishing quarantine, ensuring that the average weights of all 3 groups were as similar as possible prior to RTX-117 administration.

### CMT2N mouse model

AlaRS^R329H/+^ mice were generated using CRISPR/Cas9-mediated gene editing on a C57BL/6 background at the School of Public Health (Shenzhen), Sun Yat-sen University. Germline transmission of the specific mutation was confirmed by PCR and genomic DNA sequencing, and heterozygous lines were established. For the analysis of ISR pathway activation, spinal cord tissue was harvested from 2-month-old male and female mice. To minimize experimental variation, all analyses were performed by comparing heterozygous mice with their age-matched wild-type littermates.

### CMT2A mouse model

CMT2A mouse model work was provided as a service by WuXi AppTec (Shanghai, China). *Tg- MFN2^R94Q^* mice were obtained from the Jackson Laboratory (B6; Prp-MFN1 MFN2^R94Q^, JAX stock #033391). Genotypes were confirmed by PCR analysis of genomic DNA isolated from tail clips as previously described. For the analysis of ISR pathway activation, spinal cord tissue was harvested from 2-month-old male nTg and *Tg-MFN2^R94Q^*littermates.

### In vivo drug administration

RTX-117 was prepared as a clear solution in DMSO: 20% HPBCD in water at a ratio of 1:99 (for VWM mouse studies) or 10% (v/v) TPGS in H_2_O (For CMT2D mouse studies). Dosing solutions were prepared on the first dosing day, prepared weekly thereafter, and stored at 4°C until use. RTX-117 suspension was mixed prior to administered by oral gavage at the indicated dose level per kg body weight.

Customized chow containing two concentrations of RTX-117 (30 and 300 mg/kg) was produced by Jiangsu Xietong Pharmaceutical Bio-engineering Co., Ltd (Nanjing, China). After mixing and following routine production, chow was sterilized by irradiation. Similarly prepared chow but without added compound was administered as a vehicle diet. Vehicle or RTX-117 formulated chow was replenished weekly.

### Pharmacokinetics

Six- to eight-week-old C57BL/6 or *Gars^P278KY/+^* mice were dosed orally with a single dose of RTX- 117 at the indicated concentrations. Blood was drawn at the following timepoints post-dosing: 0.5, 1, 2, 4, 8, 12 and 24 hours and processed to plasma as described below. During the *Gars^P278KY/+^*mice study RTX-117 exposure in the plasma was determined by collecting submandibular bleed as an in-life procedure at day 2, 4, 7, and weekly thereafter or by cardiac puncture at terminal timepoints and analyzed using LC/MS/MS.

### Behavioral testing

Animals were habituated to the testing room for at least 30-60 minutes before each session. Body weight measurements: Body weight was recorded weekly throughout the study using a calibrated digital scale. Weights were recorded at the same time of day and prior to any gavage dosing on measurement days.

#### Beam walk

Mice were trained to traverse a 2.5 cm beam toward a darkened goal box starting with a distance of 10 cm away from the box, and advancing to 15cm, 30cm, 45cm, 60cm repeated twice prior to test. Mice were tested on a 2.5 cm beam at a distance of 60 cm away from the box. A video camera was set up to record animals progressing across the beam. For CMT2D mice, time to cross the beam was recorded for successful crosses. If a mouse fell, the time and traveled distance was recorded. The procedure was repeated 4 times. For VWM mice, the number of foot slips and falls were recorded.

#### Grip strength

Grip strength was measured using a digital force meter (Bioseb, BIO-GS4). Animals were held by the tail and allowed to grasp the metal grid bar with all four limbs before being pulled back at a consistent rate. Each animal performed 5-8 consecutive trials, with 1 min rest between trials. The mean of 3 trials was recorded and normalized to body weight (g force/g body weight).

#### Rotarod test

Mice were trained on an accelerating rotarod (Panlab, LE8305) for 4 consecutive days before testing at a fixed speed from 4 to 14 rpm. On the test days, the rod accelerated from 4-40 rpm over 90s. The latency to fall and speed of each mouse was recorded.

#### Electrophysiology

For all recordings, mice were anesthetized with isoflurane and maintained at a 37℃ on a heated pad throughout the experiment. Two sterile acupuncture needles (diameter, 0.25mm) were inserted into Achilles tendon (AT), and then sciatic notch (SN) and stimulated with a constant-voltage (20 v) square-wave pulses (duration, 0.1 ms; interval 3s). Compound muscle action potentials (M waves) were recorded by two acupuncture needles (diameter, 0.25mm) inserted into the interosseous muscle of hindpaw. The firings were monitored using standard electrophysiological techniques and recorded onto a PC using CED Spike 2 software (Cambridge Electronics Design). An average of 10 recordings was measured. The latencies of the compound muscle action potentials were measured and recorded. The latency from the stimulus artifact to the onset of the negative M-wave deflection or to the positive peak time of M-wave was calculated. The distance between SN and AT was measured and recorded. MNCV= (sciatic M wave latency - Achilles tendon M wave latency)/distance between SN and AT stimulation point.

### Histopathology and Immunohistochemistry

Tissue Collection and Processing: Tissues were fixed by transcardiac perfusion with saline followed by 10% neutral buffered formalin. Brains were excised and allowed to post-fix in 10% neutral formalin. Following fixation, the brain tissues were trimmed, dehydrated, embedded in paraffin, and sectioned according to standard protocols.

Staining and Imaging: For the evaluation of myelin and astrogliosis, sections were stained with Luxol Fast Blue (LFB) solution (Cat. No. L0294, Sigma-Aldrich) and an anti-GFAP antibody (Cat. No. MAB3402, Millipore), respectively. For GFAP immunohistochemistry, positive signals were visualized using a DAB substrate kit (Cat. No. BL732A, Biosharp). Following staining and mounting, the whole-slide sections were scanned and digitized using an MF43-N slide scanner (MShot).

Image Analysis and Quantification: Digital image analysis and quantification were performed using QuPath software (v0.5.1)(*67*). To evaluate the corpus callosum for both LFB and GFAP stains, specific regions of interest (ROIs) encompassing the corpus callosum were annotated on a single section at the same coronal level for each animal tissue. Within each defined ROI, the software calculated the total area and the percentage of the area exhibiting positive staining (Positive %). This positive area percentage served as the representative quantitative value for each animal. The same parameters and color deconvolution thresholds were uniformly applied to all images.

### Measurement of GDF15 and FGF21 protein levels by MSD assay

WT and CMT2D mouse CSF and plasma samples were analyzed according to manufacturer’s protocol using the following Meso Scale Discovery (MSD) kits: R-PLEX Mouse GDF-15 Assay (MSD, catalog # K152APNR-2) and U-PLEX Mouse FGF-21 Assay (MSD, catalog # K1525WK- 2). Plates were read on a MESO QuickPlex SQ 120MM (MSD) plate reader and data captured in Methodical Mind software (Version 1.0.38). Concentrations for each sample were determined by MSD analysis software using a 4-parameter logistic model. Samples from each animal were run with technical replicates and results averaged.

### Sample collection and RNA isolation

Animals were transcardially perfused with ice-cold PBS. Tissue was collected, weighed, frozen on dry ice or RNAlater reagent for samples for RNA, and stored at –80°C for subsequent analysis.

Whole blood collected into EDTA-coated tubes was spun down at 12,700 rpm for 7 min at 4°C before collecting the top plasma layer for further analysis.

Total RNA was isolated from ∼20 mg pieces of frozen tissue from brain, sciatic nerve, or spinal cord tissue using RNAzol in a tissue homogenizer System (ShanghaiJingxin-JXFSTPRP-CL) set at 60 hz for 1 min repeated 5 times with 30s pause between pulses. After tissue was completely homogenized, total RNA was isolated using VAMNE Magnetic Universal Total RNA Kit (Vazyme-ROA3301) according to manufacturer’s instructions. The CMT1X mouse model was as previously described(*53*)(*68*). Sciatic nerve samples (5-10 mg) were collected from 2-, 4-, and 6- month-old CMT1X (Cx32-null) and WT mice and stored in RNALater (ThermoFisher Scientific/Life Technologies) at - 80 C.

### Bulk RNA-seq of human PBMCs and mouse tissue samples

RNA-seq libraries were prepared from total RNA isolated from various samples as described above. For PBMC samples, mRNA was enriched using the Dynabeads™ mRNA DIRECT™ Micro Purification Kit (Thermo Fisher Scientific, 61021). For mouse brain, sciatic nerve, and spinal cord samples, ribosomal RNA was depleted using the Ribo-clean rRNA Depletion Kit Mega (Vazyme, RN416-03). Strand-specific RNA-seq libraries were generated using the VAHTS Universal V8 RNA-seq Library Prep Kit for MGI (Vazyme, NRM605-02). The resulting libraries were subsequently sequenced on an MGI T7 platform utilizing a paired-end 150-base pair (PE150) sequencing strategy by a commercial service provider. For RNA samples from Cx32-null and WT mice, RNA extraction, purification, library preparation, and sequencing was conducted by GeneWiz (Azenta Life Sciences) using the Illumina NovaSeq platform.

### Transcriptomics analysis: differential expression, gene set enrichment analysis, and single-sample pathway activity scoring

Transcript-level expression was quantified from RNA-seq reads using salmon (v1.10.0)(*69*). A transcriptome index was built using Ensembl release 112 with the GRCm39 genome used as a decoy. Gene-level estimates were then quantified using tximport (v1.34.0)(*70*) with the ‘countsFromAbundancè parameter set to ‘n’. All differential gene expression analyses were performed using limma/voom (v3.66.0)(*71*), with sample-level quality weighting(*72*).

Gene set enrichment analysis was performed downstream of differential expression analysis by limma using the fgsea package (v1.36.2)(*39*). Unless otherwise noted, genes were ranked by their t-statistics for testing. Standard pathway definitions were taken from the hallmark(*40*) and reactome(*41*) databases. The collection of ISR related signatures and their use is described in the following “ISR Signature Collection” section.

Single-sample pathway activity scores were calculated similarly as described(*73*) and implemented in the eigenWeightedMean function of the sparrow bioconductor package(*74*). Briefly, the activity score for a given sample is based on the weighted mean expression (log_2_(CPM)) of the genes in the signature; weights are assigned to genes based on their loadings on to the first principal component calculated from a principal components analysis of the expression matrix that consists of only the genes in the given signature. Significant differences between activity scores across groups was evaluated using estimated marginal means from a linear model fit with group as a categorical predictor and no intercept (lm(activity_score ∼ 0 + group [+ optional_batch_covariates])(*75*). Contrasts were computed using the emmeans R package(*76*), and p-values were adjusted across contrasts using Holm’s sequentially rejective procedure to control the family-wise error rate(*77*).

#### Assessment of ISR modulation by RTX-117

Assessment of ISR activation in a given sample was performed via GSEA using the seven primary ISR signatures outlined below. In all cases, positive normalized enrichment scores (NES) with FDR < 10% were considered to initially confirm ISR activation. Because not all genes within a primary signature are upregulated in a particular context, we created a specific ISR signature for each context to more precisely track the degree of ISR down-regulation by RTX-117 per experiment. For each experiment, the trimmed signatures were created by identifying the set of genes upregulated ( >= 30% at an FDR <= 5%) within the initial ISR-positive signature(s). The expression values of the genes within this trimmed signature were used to assess ISR modulation, either by standard GSEA or single-sample signature scores followed by linear model testing, as described above. The trimmed ISR signatures used for the PBMC/Tg, VWM, and CMT2D study are found in **Table S3B, S3D, S3F**.

#### ISR activation and RTX-117 modulation in human PBMCs

ISR activation in thapsigargin treated PBMCs was confirmed by GSEA (**Table S3A**). A thapsigargin/PBMC specific 106-gene ISR signature was generated from the genes in the ISR Class 1 signature that were upregulated > 30% at FDR < 5% (**Table S3B**). Analysis of single-sample ISR scores from the 106-gene signature was performed by linear model, with p-values corrected using Holm’s method to show significant, dose-dependent down regulation of ISR by RTX-117, which was also confirmed via GSEA analysis (**Table S3A**).

#### ISR activation and RTX-117 modulation in VWM mouse

ISR activation was confirmed by performing GSEA analysis between the untreated R191H samples vs WT controls, resulting in five of the seven ISR signatures showing significant upregulation (NES > 0, FDR < 1-6; **Table S3C**). The trimmed signature was generated from the upregulated genes in the ISR Class 1 signature (**Table S3D)**. Degree of ISR modulation was quantified using single-sample scoring from the trimmed signature, and statistical tests were performed by linear model fits as described above to show dose-dependent down regulation of ISR by RTX-117.

#### ISR activation and RTX-117 modulation in CMT2D mouse

Assessment of ISR activation and subsequent modulation by RTX-117 was accomplished using a linear model parameterized with a ‘treatment_group’ (’WT untreated’, ‘CMT2D untreated’, ‘CMT2D RTX117 lowdosè, and ‘CMT2D RTX117 highdosè) and ‘time_point’ (’2 weeks’, ‘4 weeks’, and ‘8 weeks’), like so: ‘y ∼ 0 + treatment_group + time_point’. Dysregulated pathways in CMT2D were evaluated by performing differential expression analysis between the untreated CMT2D vs WT ‘treatment_group’ followed by GSEA using signatures from Hallmark, Reactome, our primary ISR signatures (NES > 0, FDR < 10%; **Table S3E**). The CMT2D-specific trimmed ISR signature was created by extracting all genes across the six ISR-positive primary signatures that were upregulated >30% with FDR < 5%. This resulted in a trimmed ISR signature consisting of 20 genes (**Fig 3C; Table S3F**), which was used to generate single-sample ISR activity scores. RTX-117 mediated ISR modulation was tested using the same linear model parameterization. Global RTX-117 mediated rescue of other dysregulated pathways was performed via GSEA between the ‘CMT2D RTX117 highdosè and ‘CMT2D untreated’ groups, with statistics reported in **Table S3G**.

### ISR Signature Collection

A variety of gene sets have been used in the literature to show transcriptional activation of the ISR. We collated a subset of these signatures for this study, which are named “primary ISR signatures” and further “groomed” them in a manner explained below. The genset definitions derived from these analyses are included in **Table S2**.

#### Primary ISR Signatures

A. Four signatures were extracted from a previous publication(*37*) establishing a synthetic system in U2OS cells that was able to uniquely activate the ISR through specific activation *PERK*. The raw data for the class 1-3 signatures are available in GSE273601 and the ATF4 dependent signature is GSE273599:

a. **ISR Class 1 (dose responsive)**: A set of 841 genes that exhibit increased activation in a time-dependent manner with PERK activation
b. **ISR Class 2 (pulse)**: a set of 169 genes that are activated early upon PERK activation, but later return to baseline
c. **ISR Class 3 (high dose)**: a set of 205 genes that are activated after the class 2 genes have returned to baseline, and tend to be co-activated with the class 1 genes during later timepoints of ISR activation
d. **ISR ATF4 Dependent**: This was taken from Fig 3 of Labbe et al and consists of a set of genes that are upregulated during PERK activation, but not responsive during PERK activation in an ATF4/KO background.
B. **ISR CLIC Signature**: A 95 gene signature defined previously(*31*). Briefly, this signature was seeded from a core set of 50 genes that are upregulated in the cerebellum of an R191H mouse model of VWM, and further expanded to a larger set of genes that are well correlated with the core using a number of public gene expression datasets.
C. **ATF4 Targets**: A collection of genes that have been computationally inferred to be part of the ATF4 mediated regulon(*78*)(*79*).
D. **ISR SIFI**: A set of 40 ISR genes(*59*) specifically upregulated under defects of mitochondrial import.

#### Groomed Signatures

ISR responsive genes have previously been reported to be context specific(*37*), and this has also been shown by the heterogeneous upregulation of ISR genes in the PBMC/Tg, VWM, and CMT2D data presented in this study. To improve our ability to detect ISR activity in heterogeneous public RNA-seq datasets, we therefore generated context-specific versions of the primary ISR signatures by identifying the subsets of genes from each signature that are upregulated under conditions that have been previously reported to trigger the ISR.

Context specific versions of the primary ISR signatures were generated by identifying the sets of genes within each of the primary signatures that were upregulated within each of the ISR-positive experimental conditions from the experiments below:

1. ArrayExpress datasets E-MTAB-15246, E-MTAB-15247, E-MTAB-15273, E-MTAB-15295, E-MTAB-15300: Bulk RNA-seq data from different tissues was generated from an Eif2b1/N208Y mouse that has systemic activation of the ISR. Differential expression analyses were performed between Eif2b1/N208Y vs WT mice within each tissue to generate tissue specific versions of the primary ISR signatures(*80*).
2. GSE135539: RNA-seq from spinal cord of SOD1 mouse was compared vs WT controls to identify subsets of genes across primary signatures(*81*).
3. GSE233669: RNA-seq from neocortex of TDP-43/rNLS8 mice vs WT controls(*82*).
4. GSE240150: VWM Mouse (*32*).

The signatures were put into the “ISR groom” section of the ISR Signature Definition table (**Table S2**). For the exploratory analyses described in **Figure 5**, these groomed signatures were used in addition to the primary signatures in order to identify ISR-activated experimental conditions that had NES > 0 and FDR < 10%.

### Perturb-seq reanalysis and rare disease association

Reprocessed and harmonized perturb-seq data were downloaded from the *Perturbation Catalogue* web-portal (https://www.ebi.ac.uk/perturbation-catalogue/; access date May 19, 2026) hosted by the European Molecular Biology Laboratory’s European Bioinformatics Institute (EMBL-EBI; **Table S7**). The data are provided in a long format where each row contains the differential expression (DE) statistics observed from the pooled cells of each knocked out gene within each experiment. The number of differential expression statistics reported for each knocked out gene can be variable, with some KO’s only reporting statistics for just one gene, while other gene KO’s have differential expression statistics for over 4000 genes.

We performed GSEA analysis for a gene KO in a given experiment if the data reported the DE statistics for at least 250 genes. This filtered the number of genes we can test across these three datasets from 10,042 possible genes down to 475. The set of genes we could test for ISR activity was further reduced to 389 due to the overlap of the reported DE statistics with genes in related ISR signatures. Next, we performed GSEA using the ISR signature collection and the hallmark gene signatures, with genes ranked by their reported log_2_ fold change. A signature was considered to be significantly regulated if it had an FDR < 10%. Finally, to aggregate the GSEA results for each gene from the individual experiments, we summarized the normalized enrichment score (NES) for each gene and pathway combination by reporting the mean NES for the pathway among all of the individual results with FDR < 10%. These mean NES values are visualized in the heatmap shown in **Fig 5B**, with the NES statistics provided in **Table S5C**. If a particular pathway showed no significant activity in any experiment for a particular gene, no NES is reported and is visualized as a white tile in the heatmap. To keep the heatmap concise, pathways that showed significant activity across less than 10 genes are only reported in the table but not visualized in the heatmap.

Genes were grouped into functional categories by performing an overrepresentation analysis (fisher’s exact test) from gene ontology (GO) and Reactome pathways associated with the 108 genes ISR-activating genes using the 475 genes as the universe. The reactome and GO signatures that were grouped into higher-level functional categories are listed in **Table S5B**.

We used the data made available by *The Gene Curation Coalition* (GenCC) to associate genes with potential diseases of interest. Briefly, GenCC provides a curated and standardized table with associations between gene mutations and disease. These associations come with evidence codes that classify the strength of gene-to-disease association (definitive, strong, moderate, etc) as well as the inheritance pattern (autosomal dominant, autosomal recessive, X-linked, etc). Importantly, it also provides annotations that refute previously reported gene-to-disease associations. We downloaded the data from GenCC (May, 2026) and only considered gene to disease associations that were classified as either “definitive” or “strong” evidence. Any gene that has “refuted” lines of evidence for disease association were removed from consideration.

## Statistical analysis

Data are reported as means ± SD as indicated in figures legends. Statistical analysis of data was performed in GraphPad Prism version 10 or later. Analysis of data from animal studies was performed using t test or one-way analysis of variance (ANOVA) with multiple comparison (Bonferroni’s Multiple Comparison Test).

## List of Supplementary Materials

Figure S1 to S13

Table S1 to S7

Movie S1 to S4

## Acknowledgments

We thank Dr. Manfu Wang and Dr. Guoqiang Li from Biortus Discovery Co., Ltd for help with structural studies and the WuXi AppTec Neuroscience and Rare Disease Pharmacology Team, especially Dr. You Qi for help overseeing the CMT2D mouse model study. Thanks to Xtalpi team members including Liyu Zhao, Peipei Chen, Dan Yin, Ling Zhao, and Dewei Xie for experimental support. We thank Nicholas Mann for managing the RTX-117 program, Dr. Griff Humphreys for help with DMPK studies and interpretation, and members of ReviR Therapeutics Inc. for program feedback. Thanks to Dr. Sarah Burnett for critical reading of the manuscript.

## Author contributions

Conceptualization: M.M., S.L., S.N., Y.Li., P.Y., and P.R.A.

Conceptualization CMT1X mouse model study: C.K.A., M.M.F., and D.G.

Data curation: M.M., S.L., S.N., and J.Z.

Formal analysis: M.M., S.L., S.N., J.Z., S.C., J.L., B.C., J.Luo., J.C., X.Z., S.Lu., X.M., L.S., A.C.N., C.K.A., M.M.F., and D.G.

Funding acquisition: Y.Li., P.Y., and P.R.A.

Investigation: M.M., S.L., S.N., J.Z., S.C., J.L., W.K., Y.L., C.K.A., M.M.F., and D.G.

Visualization: M.M., S.L., S.N., and J.Z.

Writing: M.M., M.D., and P.R.A. with input from all authors.

## Data and materials availability

Data associated with this study is presented in the main test or in the Supplementary material. RNA-seq data from in vitro and in vivo models will be deposited at the NCBI GEO repository (ongoing). Cryo-EM data will be deposited at RCSB Protein Data Bank (ongoing).

## SUPPLEMENTAL FIGURES

**Figure S1.**
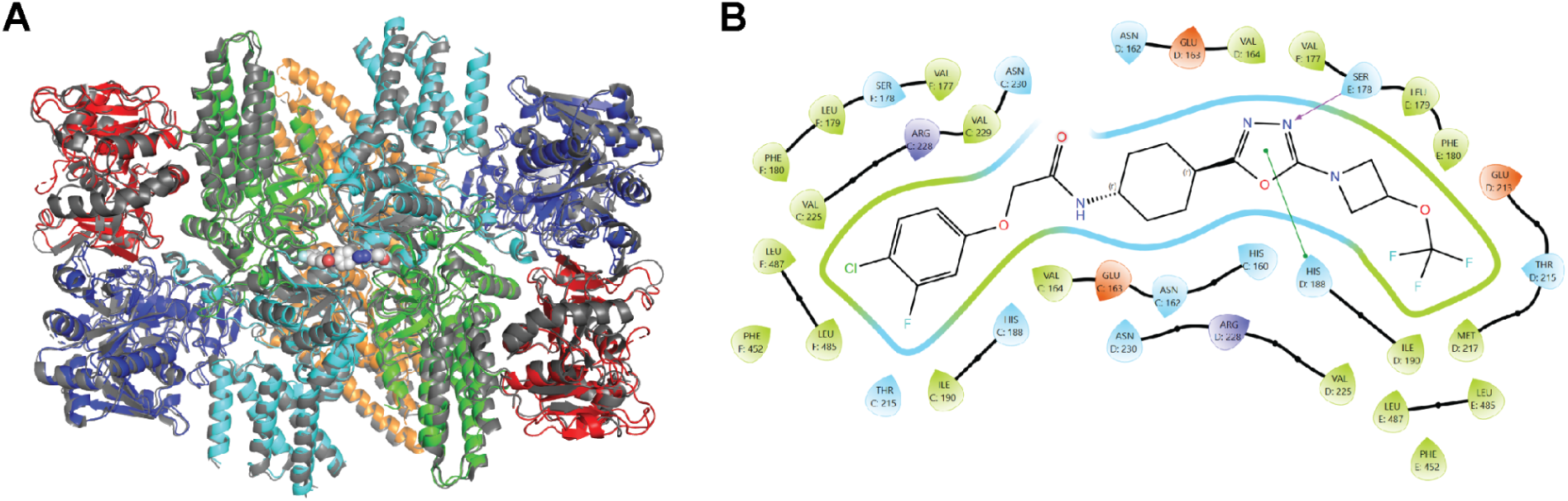
Structural analysis for RTX-117 bound to human eIF2B. (A) Overlay of the RTX-117 bound structure of eIF2B (colored) with the apo eIF2B decamer (PDB: 7L70, gray). (B) Two-dimensional ligand interaction map of RTX-117 within the eIF2B binding pocket, highlighting key hydrophobic contacts, polar interactions, and π-stacking interactions.

**Figure S2.**
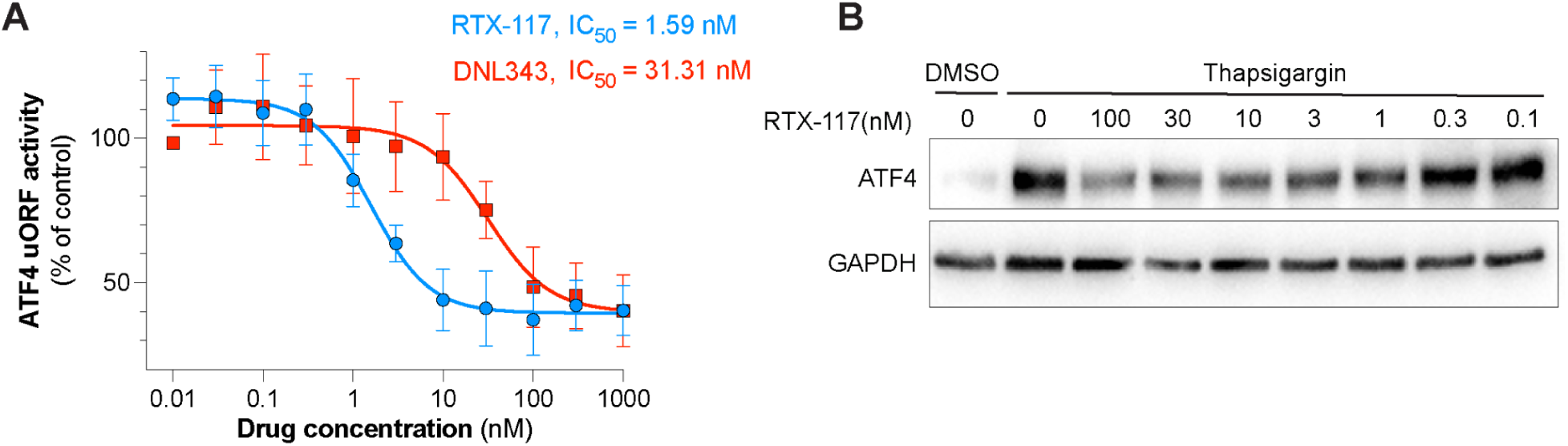
RTX-117 is an eIF2B activator demonstrating superior potency to DNL343. (A) Concentration-dependent inhibition of ATF4-luciferase reporter activity in HEK293T cells stimulated with thapsigargin. Cells were treated with increasing concentrations of RTX-117 or DNL343, and ATF4 translation was quantified using a reporter assay (data from n=5 independent experiments), Data for RTX-117 are the same as shown in Fig. 2A. Data are presented as mean ± SD. Nonlinear regression analysis yielded IC_50_ values of 1.59 nM for RTX-117 and 31.31 nM for DNL343. (B) Representative immunoblot showing endogenous ATF4 protein levels in HEK293T cells following thapsigargin treatment and increasing concentrations of RTX-117. GAPDH is shown as a loading control.

**Figure S3.**
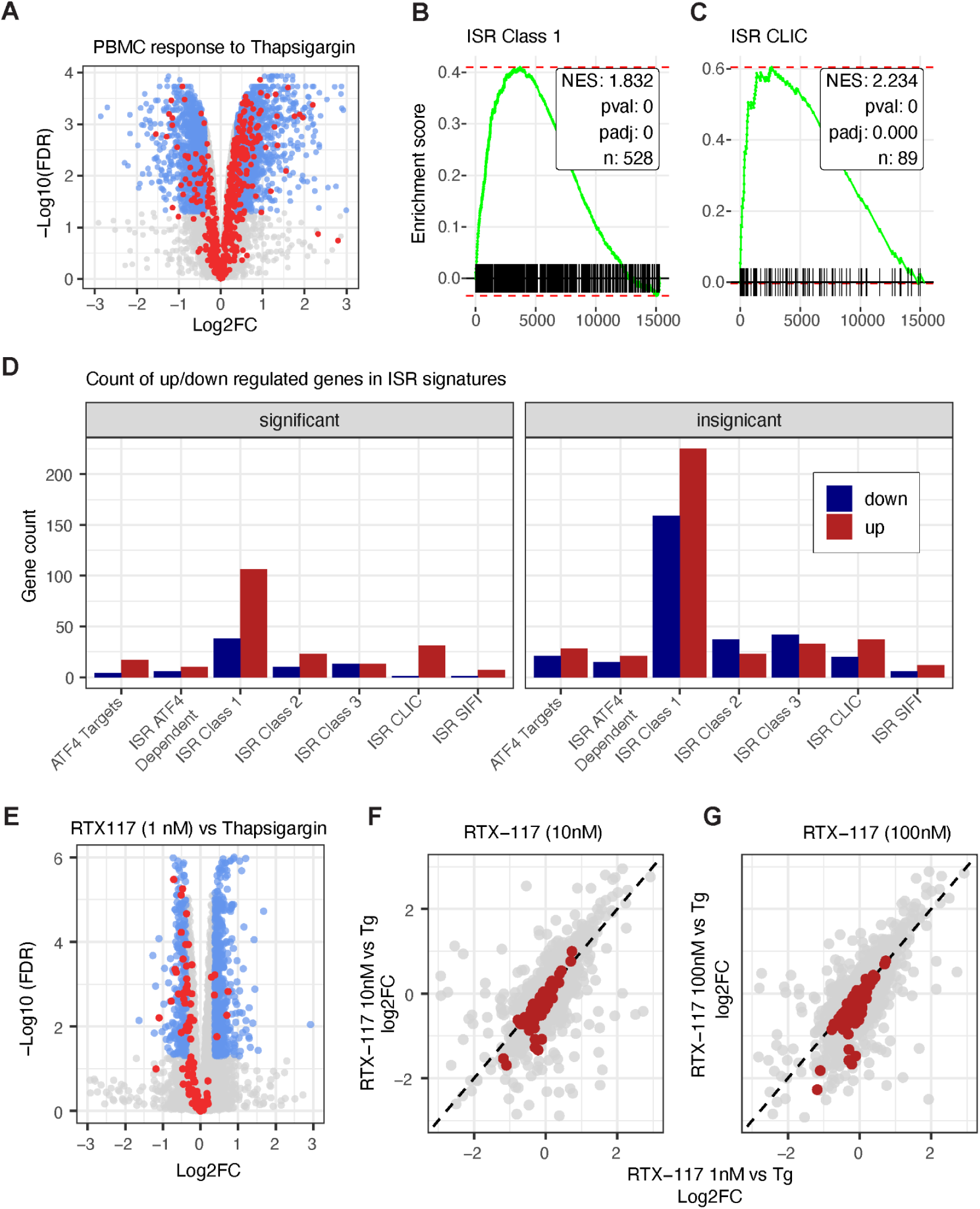
PBMC transcriptome response to thapsigargin and normalization by RTX-117 treatment. (A) Volcano plot showing differential gene expression in peripheral blood mononuclear cells (PBMCs) treated with thapsigargin. Blue dots indicate genes that are differentially expressed > 30% (up/down) at an FDR <= 5%. Red dots indicate genes from the ISR Class 1 signature. (B) GSEA results from the ISR Class 1 signature verifies significant activation of ISR. (C) GSEA results from the ISR CLIC signature verifies significant activation of ISR. (D) Number of genes that are up/down regulated across the seven primary ISR signatures from the thapsigargin treatment; genes with >30% modulation at FDR < 5% on left, others on right. (E) Volcano plots from thapsigargin treated samples co-treated with 1 nM RTX-117. Blue dots are as described in A. Red dots indicate the 106 genes from the ISR Class 1 signature upregulated under thapsigargin treatment alone. (F) Comparison of log_2_ FC of genes with 10 nM RTX-117 vs Tg (y-axis) as compared to 1 nM RTX-117 dose. Red dots are the same as in E. (G) Comparison of log_2_ FC of genes with 100 nM RTX-117 vs Tg (y-axis) as compared to 1 nM RTX-117 dose. Red dots are the same as in E. Rightward/downshift of red dots with higher doses indicates increased ISR attenuation.

**Figure S4.**
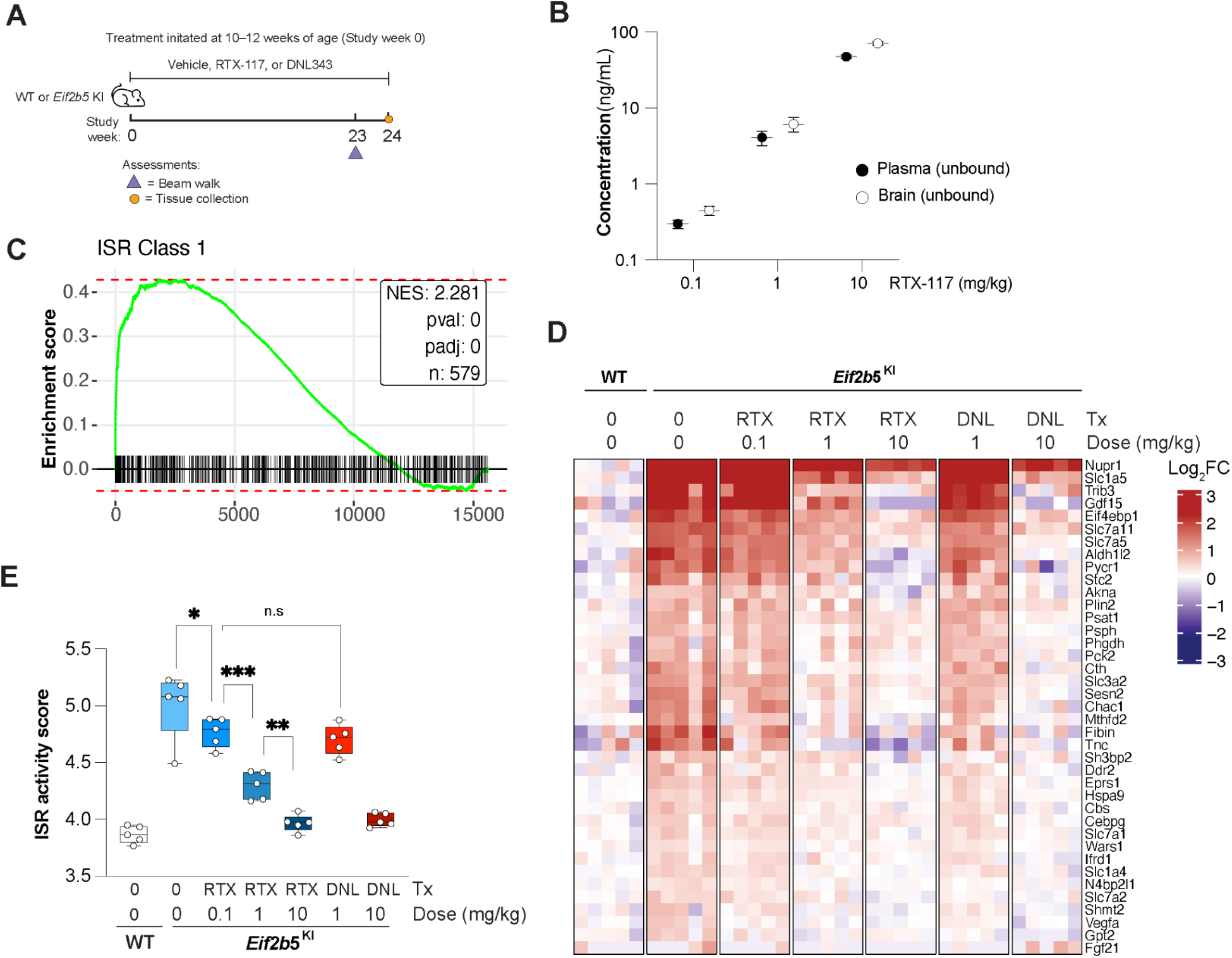
RTX-117 normalizes ISR overactivation in the brain of a mouse model of VWM disease. (A) Experimental timeline and study design. Wildtype (WT) and *Eif2b5*^R191H/R191H^ knock-in mice (Eif2b5 KI) were assigned to treatment with vehicle, RTX-117, or DNL343 beginning at 10–12 weeks of age (study week 0). (B) Unbound RTX-117 concentrations measured in plasma and brain across RTX-117 treatment groups, demonstrating central nervous system exposure (n=3). Data are presented as mean ± SD. (C) GSEA analysis shows significant upregulation of ISR Class 1 signature in untreated VWM vs WT controls (D) Heatmap of 37-gene ISR-associated gene expression in brain tissue from WT and Eif2b5 KI mice treated with vehicle, RTX-117, or DNL343 at the indicated doses (mg/kg). Expression values are shown as log_2_ fold change relative to WT vehicle treated animals from both male and female mice. The color scale represents log_2_ fold change, with red indicating increased and blue indicating decreased expression. Columns represent individual samples. Treatment groups are indicated above the heatmap (Vehicle, RTX-117 at 0.1, 1, and 10 mg/kg, and DNL343 at 1 and 10 mg/kg). (E) ISR activity scores across treatment groups is shown. Points represent individual animals and boxes indicate median and interquartile range. Stars indicate statistical significance from Holm’s adjusted p-values (see methods). * P < 0.05; ** P < 0.01; *** P < 0.001; n.s. not significant.

**Figure S5.**
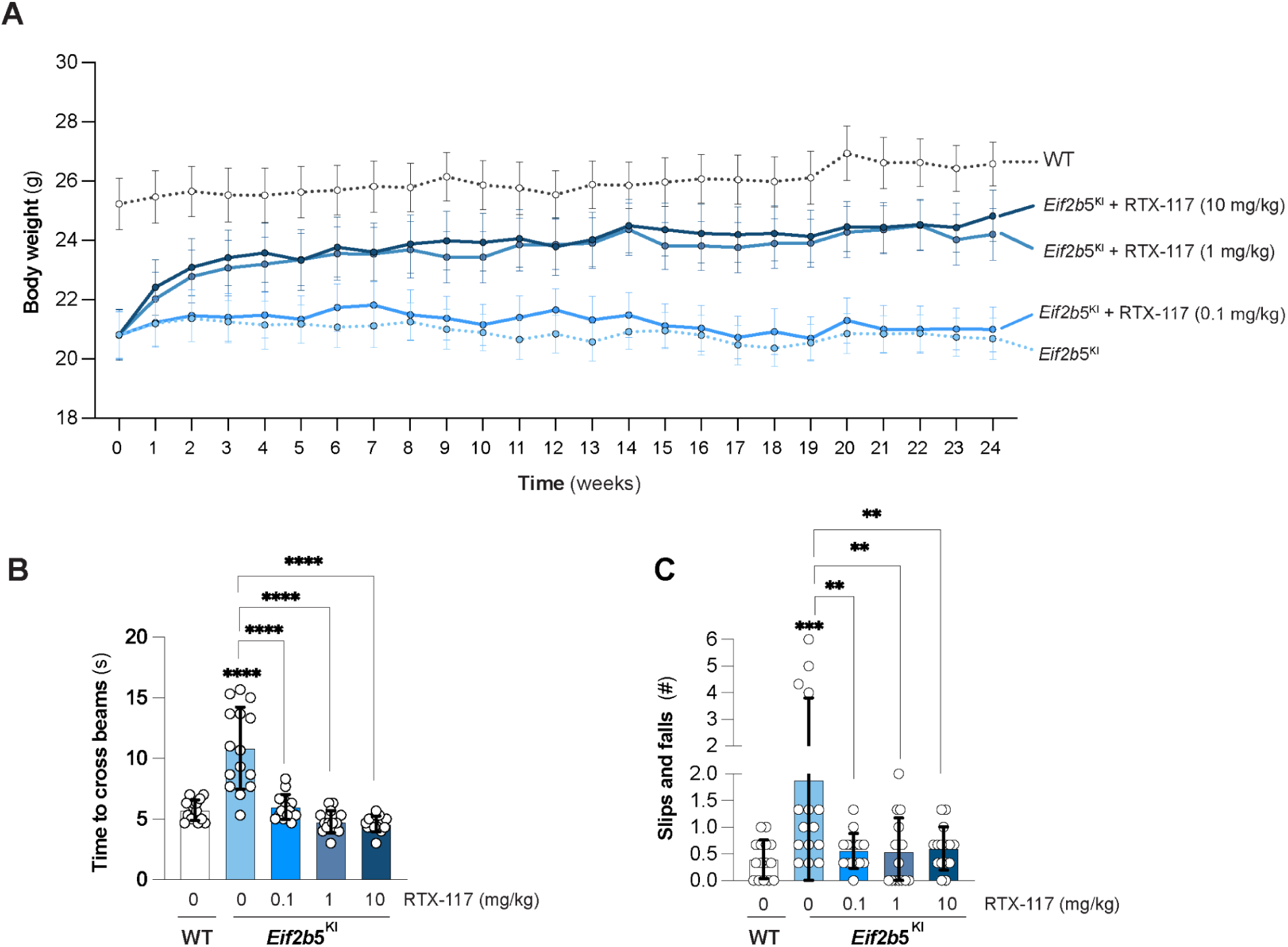
RTX-117 treatment improves body weight and motor function in a mouse model of VWM disease. (A) Body weight trajectories of male and female wild type (WT) and *Eif2b5*^R191H/R191H^ knock-in mice (Eif2b5 KI) during longitudinal treatment with RTX-117 or vehicle. KI mice received a once daily dose of RTX-117 at 0.1, 1, or 10 mg/kg or vehicle, while WT mice received vehicle treatment. Treatment was initiated at 10–12 weeks of age (study week 0) and animals were monitored throughout the study. (B) WT and Eif2b5 KI mice from all treatment groups were assessed for motor function in beam walk performance measured as time required to cross the beam. Data shown from males and females from 23 weeks post-treatment. (C) Graph shows quantification of slips and falls for mice mentioned in (B) during beam walk testing, reflecting motor coordination deficits. Data from males and females presented as mean ± SD. Statistical comparisons were performed using one-way ANOVA followed by Bonferroni’s multiple comparisons test. The KI vehicle was compared to the WT vehicle and each RTX-117 treatment group, and between the RTX-117 dose groups. Brackets denote comparisons between KI vehicles and RTX-117 treatment groups. Significance for the WT versus KI vehicle comparison is indicated above the KI vehicle bar. **P < 0.01, ***P < 0.001, ****P < 0.0001.

**Figure S6.**
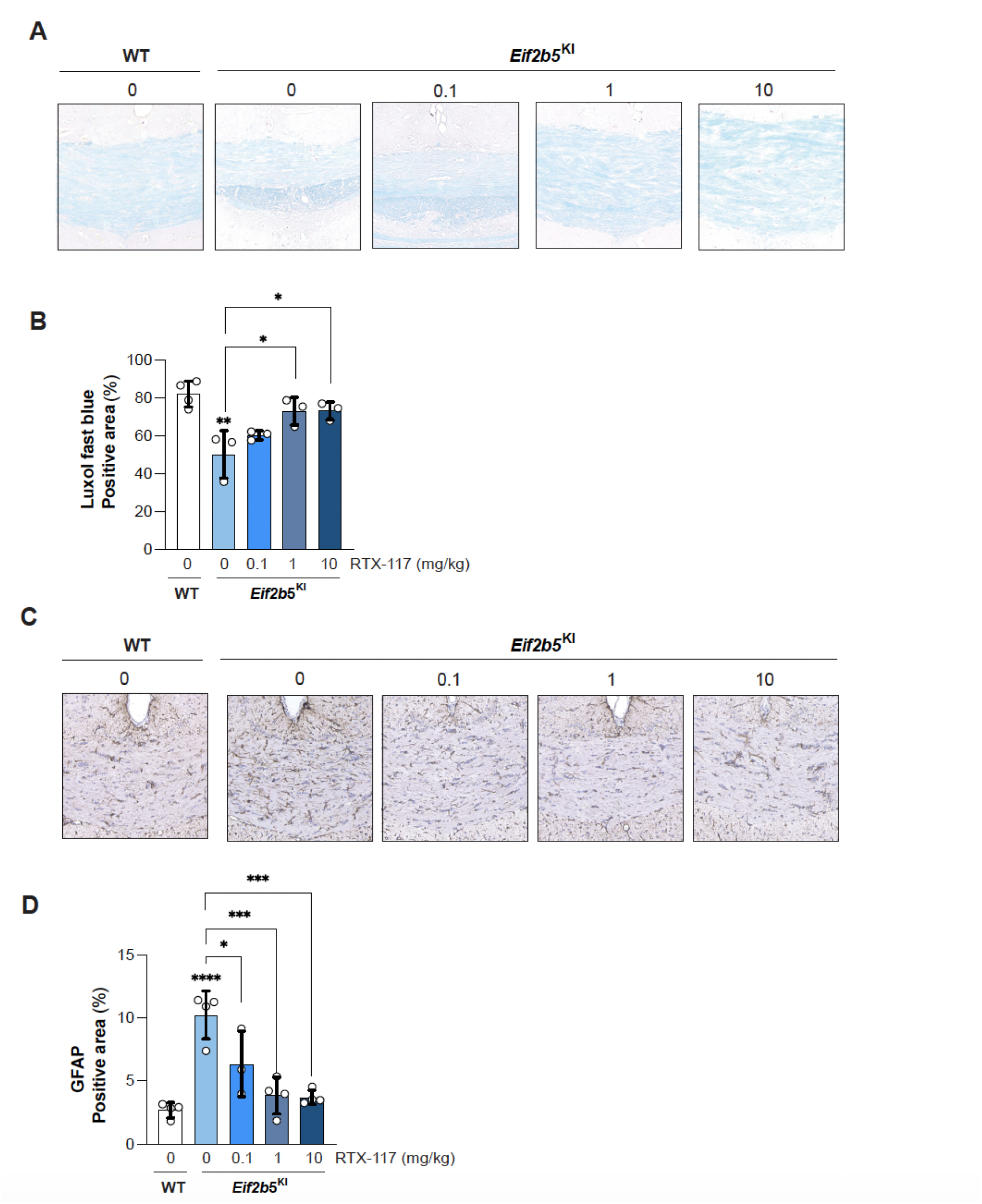
RTX-117 treatment improves neuropathological outcomes in a mouse model of VWM disease. (A) Representative Luxol fast blue staining of brain sections showing myelin integrity across WT and Eif2b5 KI mice treated once daily with vehicle or RTX-117 (0.1, 1, or 10 mg/kg). (B) Graph shows quantification of Luxol fast blue-positive area (%) across treatment groups described in A, indicating improved myelin preservation with RTX-117 treatment from both male and female mice. (C) Representative immunohistochemical staining of glial fibrillary acidic protein (GFAP) in brain sections from WT and Eif2b5 KI mice treated with vehicle or RTX-117 (0.1, 1, or 10 mg/kg), illustrating reactive gliosis. (D) Graph shows quantification of GFAP-positive area (%) across treatment groups. KI vehicle-treated mice exhibit increased gliosis relative to WT controls, which is reduced with RTX-117 treatment from both male and female mice. Data are presented as mean + SD. Statistical comparisons were performed using one-way ANOVA followed by Bonferroni’s multiple comparisons test. The KI vehicle was compared to the WT vehicle and to each RTX-117 treatment group, and RTX-117 dose groups were compared with each other. Brackets denote comparisons between KI vehicles and RTX-117 groups. Significance for the WT versus KI vehicle comparison is indicated above the KI vehicle bar. *P < 0.05, ***P < 0.001, ****P < 0.0001.

**Figure S7.**
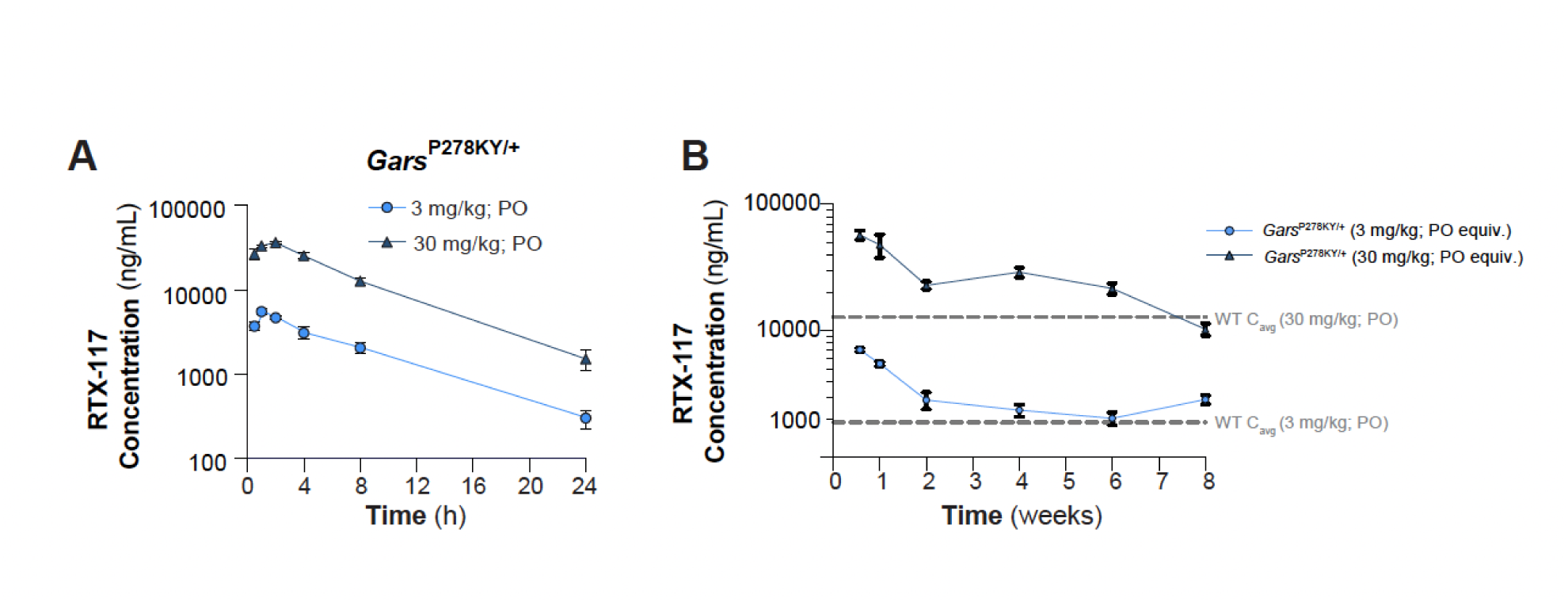
Dose-dependent plasma exposure of RTX-117 in a CMT2D mouse model. (A) Plasma PK profiles of RTX-117 measured following administration of a single PO dose in *Gars*^P278KY/+^ mice. Plasma samples were collected at indicated times post dosing and compound concentrations were quantified using LC-MS/MS. (B) Plasma PK of RTX-117 in *Gars*^P278KY/+^ mice fed RTX-117 formulated chow over a 8 week period. Dashed lines indicate mean steady-state exposure (C_avg_) observed in WT mice at 3 and 30 mg/kg PO dose.

**Figure S8.**
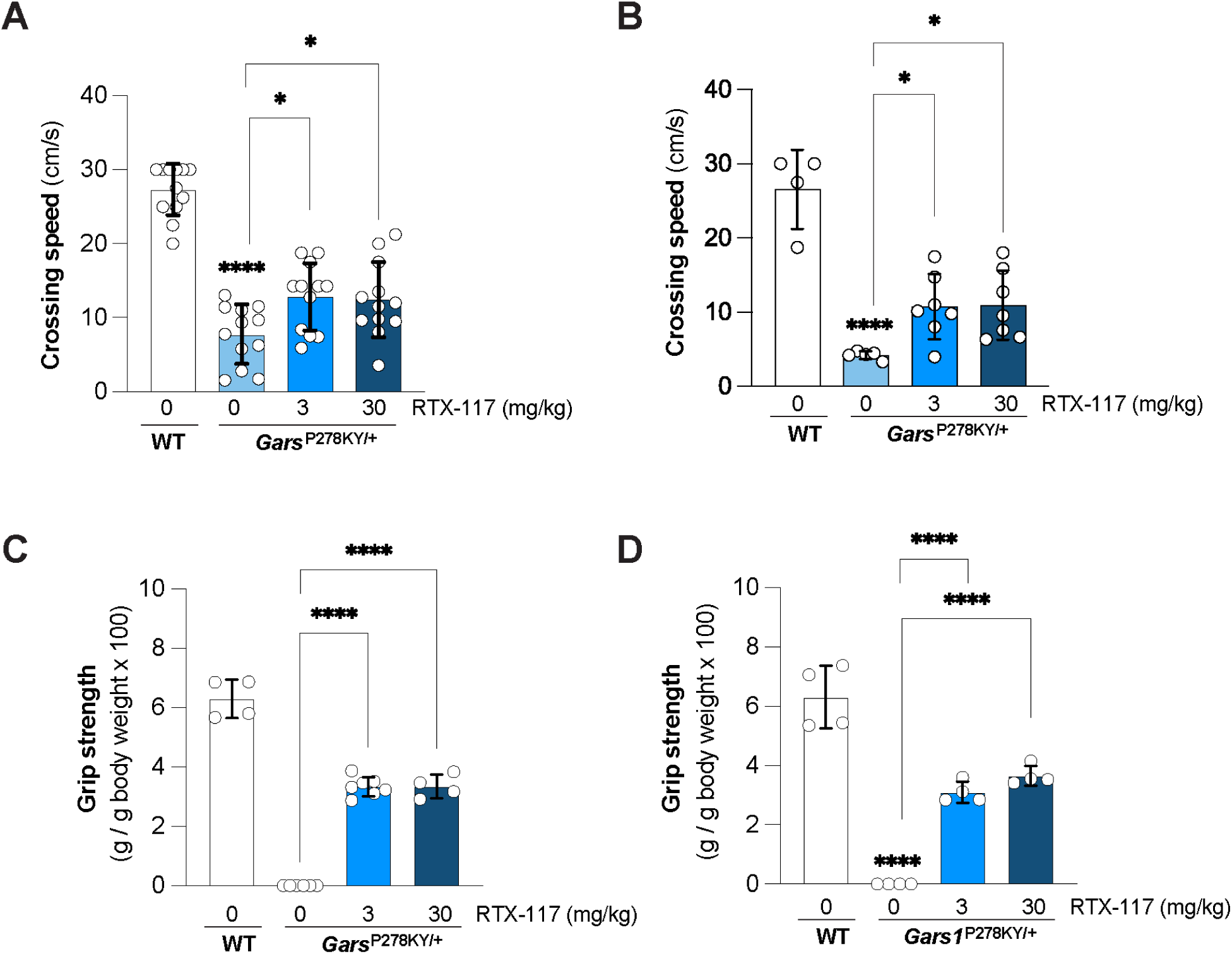
Improved neuromuscular function with eIF2B activation treatment in a mouse model of CMT2D. (A–B) Beam walk crossing speed in wild type (WT) and *Gars*^P278KY/+^ mice measured at 2 (A) and 6 (B) weeks of treatment. (C-D) Grip strength normalized to body weight in wild type (WT) and *Gars*^P278KY/+^ mice measured at 6 (C) and 8 (D) weeks of treatment. Data from males and females are presented as mean ± SD. Statistical comparisons were performed using one-way ANOVA followed by Bonferroni’s multiple comparisons test. Untreated *Gars*^P278KY/+^ mice were compared to untreated WT mice and each RTX-117-treated group, and between the two RTX-117 dose groups. Brackets denote comparisons between *Gars*^P278KY/+^ groups. Significance for the WT versus *Gars*^P278KY/+^ comparison is indicated above the *Gars*^P278KY/+^ bar. *P < 0.05, ****P < 0.0001.

**Figure S9.**
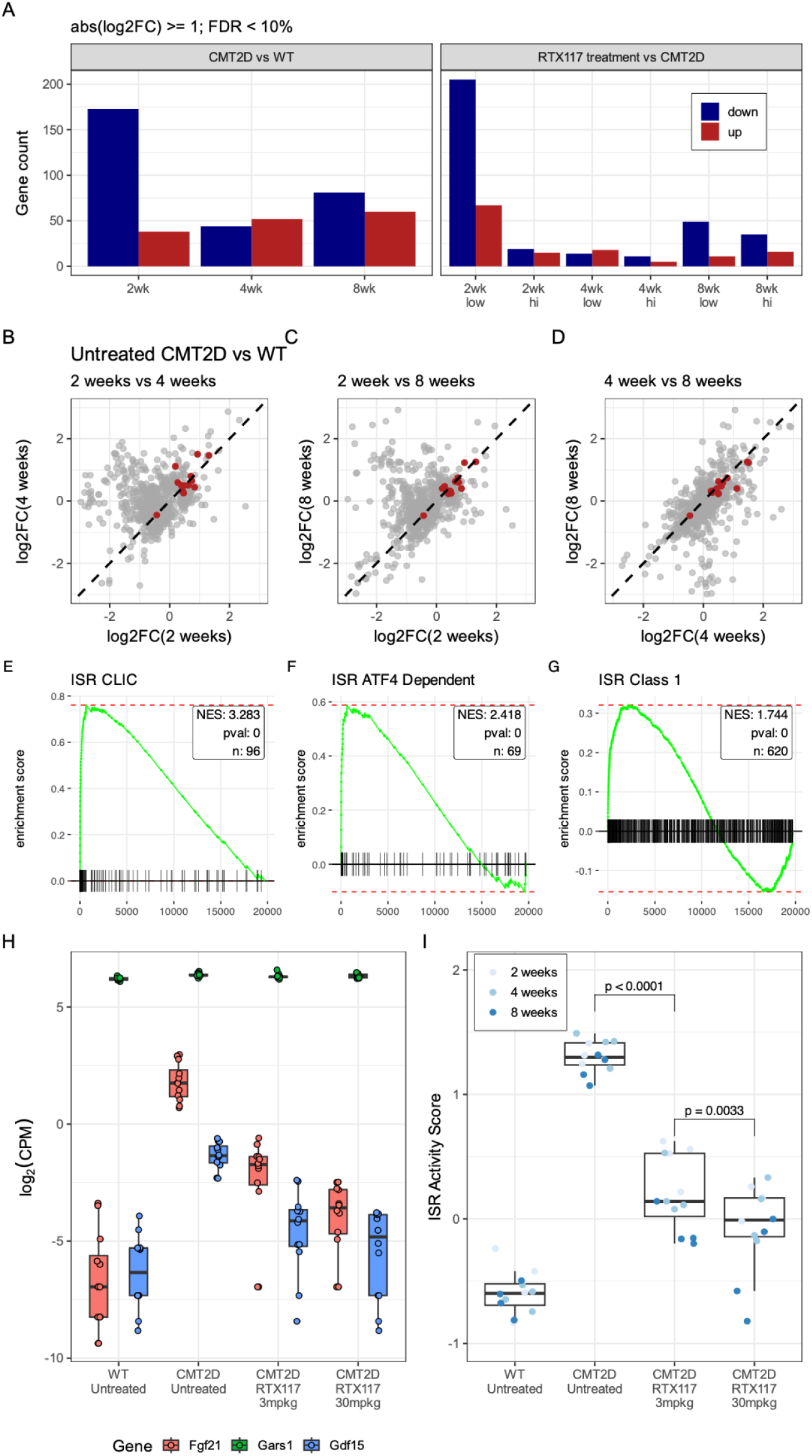
ISR activation in CMT2D spinal cord, and dose-dependent reduction by RTX-117. (A) Number of genes differentially expressed across untreated (left) and treated (right) mice. Counts are from genes up/down regulated by more than 2x at FDR <= 10%. (B-D) log_2_ FC between untreated CMT2D and WT are shown across timepoints. Genes with >= 30% up/down regulation at FDR <= 5% shown. Genes from ISR CLIC signature (depicted in red) show comparably similar levels of ISR activation across all timepoints. (E-G) GSEA results from untreated CMT2D mice vs WT show ISR activation across the majority of primary ISR signatures. (H) Expression levels of *Fgf21 (red)*, *Gars1 (green)*, and *Gdf15 (blue)* in all of the samples from the indicated groups in the study. (I) Single-sample ISR activity scores from all samples in the indicated groups in the study, colored by timepoint. Both low and high RTX-117 treated mice show statistically significant reduction of ISR activity when compared to untreated CMT2D mice. High dose RTX-117 shows statistically significant reduction of ISR activity when compared to low dose. Holm-adjusted p-values shown between comparisons.

**Figure S10.**
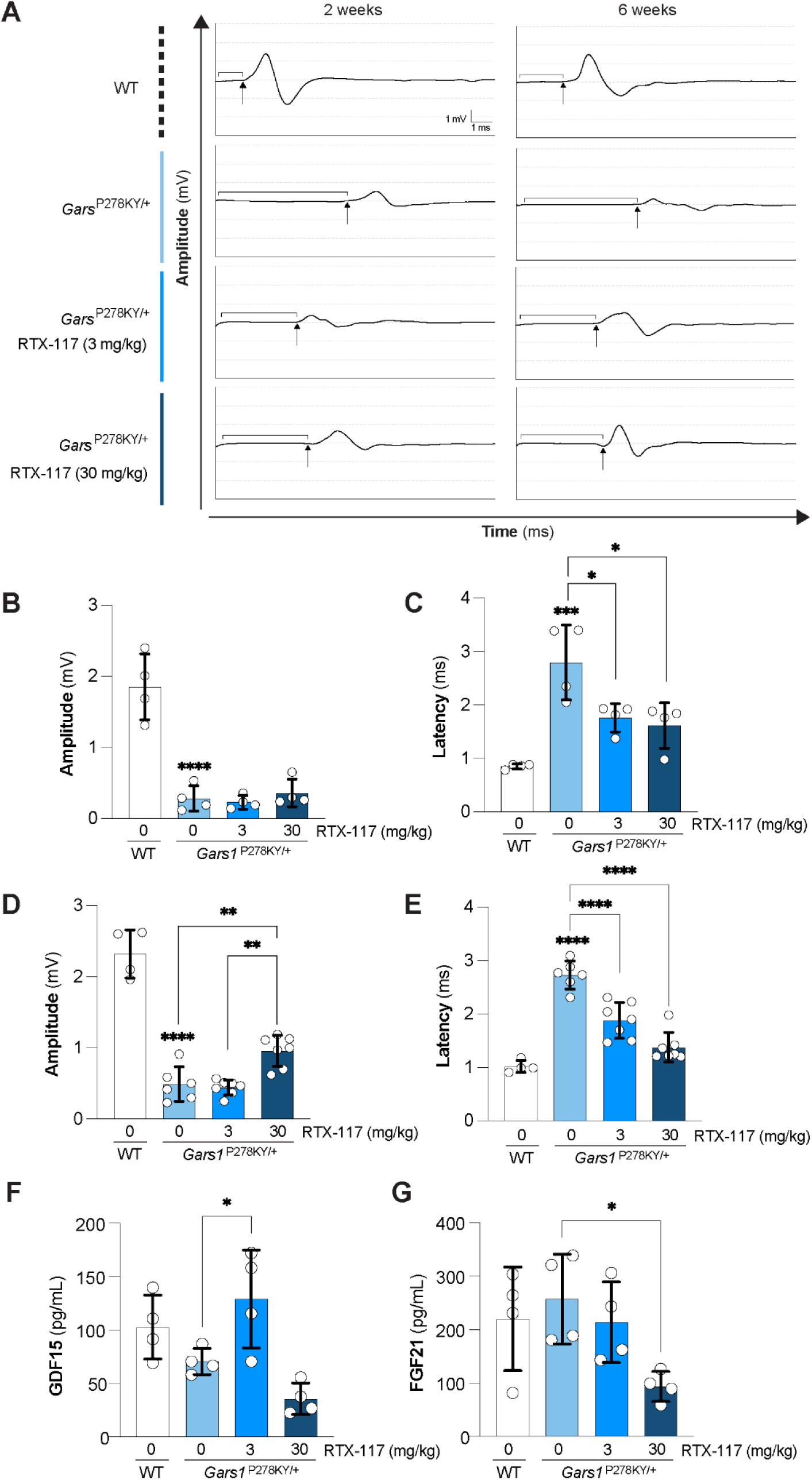
RTX-117 improves proximal CMAP in a mouse model of CMT2D. **(A)** Representative nerve stimulation evoked responses showing proximal compound muscle action potential (CMAP) recordings from wild-type (WT) and *Gars*^P278KY/+^ mice at 2 (left) and 6 (right) weeks of treatment following stimulation at the sciatic notch. Scale bars, 1 mV and 1 ms. Bracket denotes latency; arrow denotes onset. **(B–C)** Quantification of CMAP amplitude (B) and latency (C) at 2 weeks across treatment groups. **(D–E)** Quantification of CMAP amplitude (D) and latency (E) at 6 weeks across treatment groups. Data from males and females are presented as mean ± SD. Statistical comparisons were performed using one-way ANOVA followed by Bonferroni’s multiple comparisons test. (F) GDF15 protein levels in plasma from wild type (WT) and *Gars*^P278KY/+^ mice following 8 weeks of RTX-117 treatment (G) FGF21 protein levels in plasma from wild type (WT) and *Gars*^P278KY/+^ mice following 8 weeks of RTX-117 treatment. Comparisons were made between untreated *Gars*^P278KY/+^ mice and untreated WT mice, between untreated *Gars*^P278KY/+^ mice and each RTX-117-treated group, and between the two RTX-117 dose groups. Brackets denote pairwise comparisons between *Gars*^P278KY/+^ groups. Significance for the WT versus *Gars*^P278KY/+^ vehicle comparison is indicated above the *Gars*^P278KY/+^ vehicle bar. *P < 0.05, **P < 0.01, ***P < 0.001, ****P < 0.0001.

**Figure S11.**
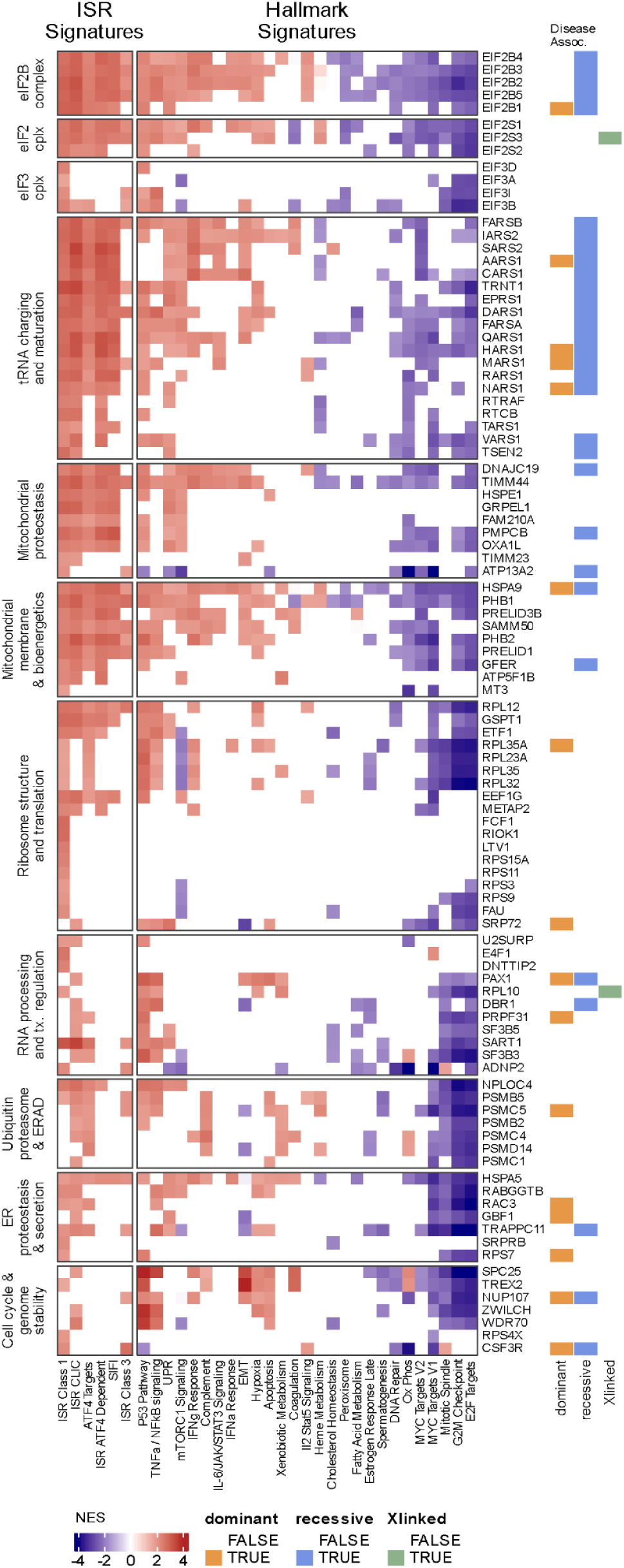
Extended Pathway dysregulation inferred from Perturb-seq data. Tiles show the mean normalized enrichment score (NES) for each gene–pathway association across cellular contexts in which GSEA reached FDR < 10%. White tiles indicate that the pathway was not significantly altered in any analyzed cellular context. Genes are organized into functional groups derived from Gene Ontology enrichment analysis. GenCC annotations at right indicate genes with strong or definitive evidence of disease association, categorized by reported mode of inheritance: dominant (orange), recessive (blue), or X-linked (green).

**Figure S12.**
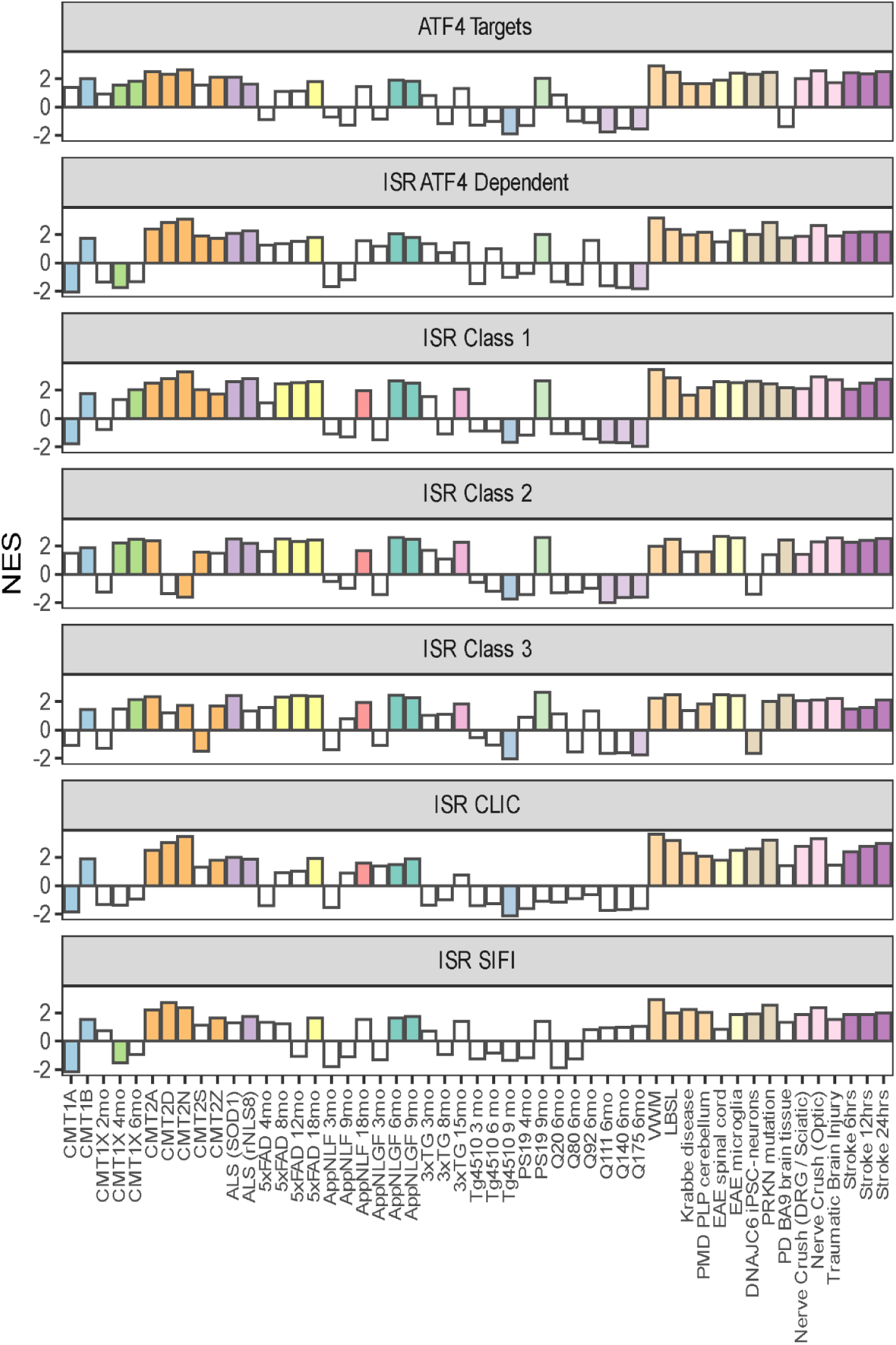
Extended GSEA analysis for neurodegenerative indications. GSEA statistics are presented from all seven primary ISR signatures. Bar height indicates normalised enrichment score (NES) from GSEA analysis of the denoted ISR signatures. Colored bars denote results meeting FDR < 10%.

**Figure S13.**
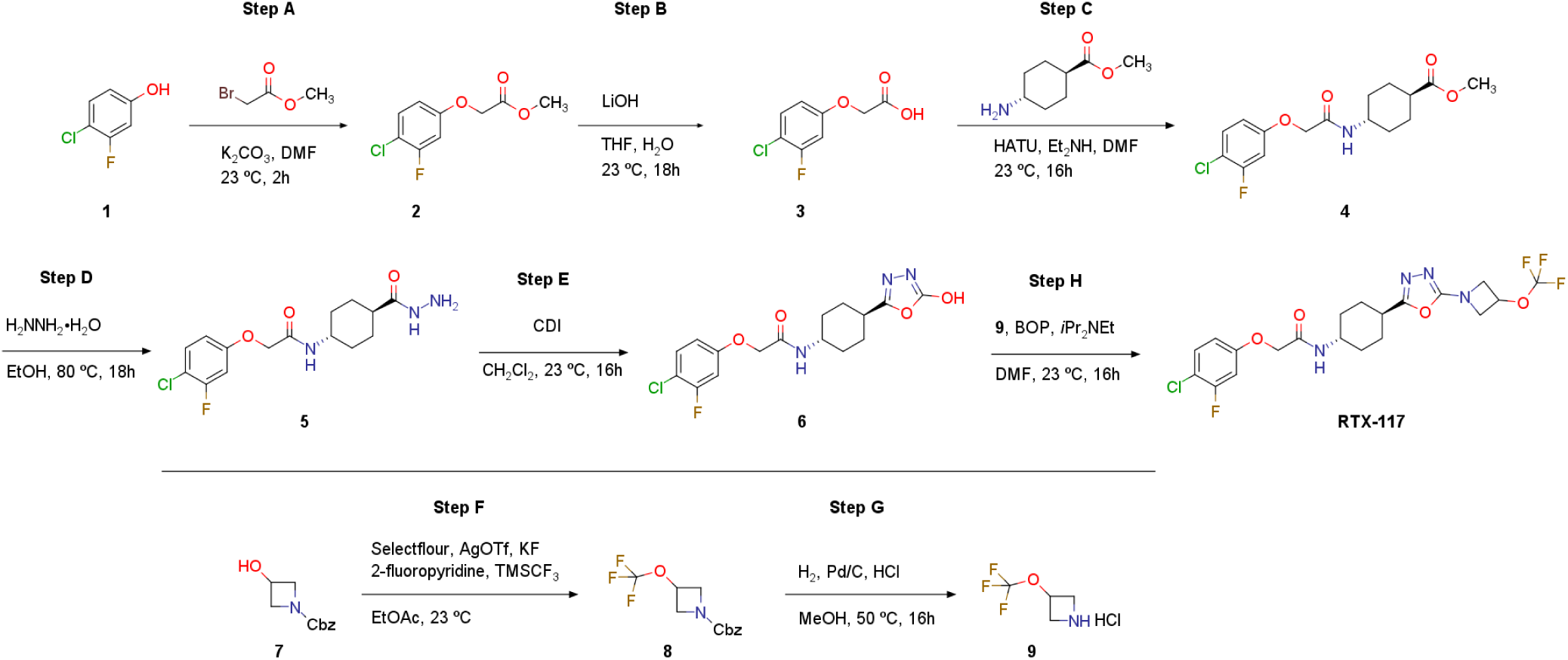
Synthetic route to RTX-117. Eight-step synthesis of RTX-117, with reagents and conditions shown at each arrow. Steps A to E build the aryloxyacetamide–oxadiazolone core (**6**); steps F and G, shown below the line, prepare the 3-(trifluoromethoxy)azetidine fragment (**9**), which is coupled to **6** in step H to give RTX-117. Isolated yields, experimental procedures, and characterization data (ESI-MS and ¹H NMR) are provided in Materials and Methods.

