## Supplementary material for "Pharmacologic eIF2B Activation Rescues Neuropathy in CMT2 Subtypes by Normalizing the Integrated Stress Response": Table S1

**Table S1. Calculated and measured physico-chemical and ADME properties of RTX-117**

| **Parameter** | **Value** |
| --- | --- |
| Mol. Wt. | 493 |
| clogP | 4.8 |
| logD_7.4_ | 4.1 |
| TPSA | 90 Å^2^ |
| H-bond donors | 1 |
| H-bond acceptors | 8 |
| CNS MPO score | 3.1 |
| BBB score | 3.14 |
| Solubility at pH 7.4 | 11.3 µg/mL |
| Caco-2 Permeability  A-B  B-A  Efflux ratio | 26  18  0.71 |
| t_½_ (human liver microsomes) | > 185 min |
| Kp,uu (mouse) | 1.4 |
